# Exploiting the polypharmacology of alectinib for synergistic RNA splicing disruption with RBM39 degraders

**DOI:** 10.1101/2025.03.28.644532

**Authors:** Yurui Ma, Evon Poon, Chenchen Jin, Barbara Martins da Costa, Yuewei Xu, Sadiya Quazi, Nikolaos Zourdoumis, Chiharu Wickremesinghe, Louis Chesler, Hector C Keun, Anke Nijhuis

## Abstract

Precise control of pre-mRNA splicing is critical for transcriptome integrity, and its disruption is increasingly recognised as a vulnerability in cancer. Here, we identify a functional interplay between two key splicing regulators, RBM39 and serine/arginine protein kinase 1 (SRPK1), and show that dual targeting of these factors severely compromises splicing fidelity in high-risk neuroblastoma. We use the molecular glue indisulam to degrade RBM39 and repurpose the clinical ALK inhibitor alectinib which potently inhibits SRPK1. Co-treatment with indisulam and alectinib inhibited cell proliferation, induced apoptosis, and caused G2/M arrest in multiple cancer cell lines, including *MYCN*-amplified neuroblastoma. RNA sequencing revealed enhanced splicing defects preferentially in DNA repair and genome maintenance related genes following combination treatment, leading to R-loop accumulation and increased DNA damage. In the Th-MYCN/ALK^F1174L^ neuroblastoma mouse model, combination therapy induced complete tumour regression and significantly improved survival rates compared with monotherapies. These findings demonstrate that combining indisulam and alectinib is a promising approach to treat aggressive malignancies such as high-risk neuroblastoma, exploiting the previously untapped polypharmacology of alectinib as a clinical RNA splicing inhibitor and supporting the therapeutic value of co-targeting interdependent splicing factors for synergistic benefit.

## INTRODUCTION

The splicing of precursor messenger RNAs (pre-mRNAs) is a critical regulatory step in gene expression, during which non-coding introns are excised to produce functional mRNA(1). This process is tightly controlled by splicing factors, governing the recognition and selection of splice sites, thereby generating mRNAs with varying exon combinations and consequently diverse protein isoforms(1). Dysregulated alternative splicing can promote tumorigenesis and chemoresistance. Aryl sulfonamides such as indisulam have emerged as promising anticancer agents by targeting an RNA splicing factor RBM39 (RNA binding motif 39)(2,3). These molecular glues facilitate the association of RBM39 with the CUL4-DCAF15 ubiquitin ligase, resulting in RBM39 polyubiquitination and subsequent proteasomal degradation(2,4). Suppressing RBM39 efficiently inhibits tumour growth and advancement through modulating the transcription of various genes associated with tumorigenesis (reviewed in (3)).

One cancer type particularly vulnerable to splicing perturbation is neuroblastoma(5), a common and aggressive extracranial solid tumour in young children, and a major contributor to paediatric cancer mortality(6). High-risk neuroblastoma, comprising approximately 15% of all neuroblastoma cases, represents a distinct subtype characterized by aggressive tumour behaviour and an unfavourable prognosis(7). One pivotal factor contributing to the progression of high-risk neuroblastoma is the amplification of the MYC family transcription factor, MYCN, which can block the differentiation of neural progenitors, the key event in neuroblastoma initiation(8), and plays a crucial role in facilitating the transcriptional repression of tumour suppressor genes(6,7). Furthermore, oncogenic gain-of-function mutations in the *ALK* (Anaplastic Lymphoma Kinase) occur in 10–15% of primary tumours, making ALK the most common single-gene alteration in high-risk neuroblastoma for which targeted clinical therapies have been developed(6,9) Alectinib, an FDA-approved ATP-competitive ALK tyrosine kinase inhibitor (TKI),was initially developed for non-small-cell lung cancer and has shown considerable promise in addressing neuroblastoma in extensive preclinical studies(10–12). However, tumour heterogeneity and resistance to treatment with ALK inhibitors (and molecularly-targeted drugs in general) limit their efficacy as monotherapies(13). Thus, novel strategies, particularly combination therapies, remain important as a means to reduce the toxicity of any single drug and overcome treatment resistance more efficiently. This approach is particularly advantageous in childhood cancer, where long-term toxicity is a major lifelong issue (14).

Our previous study(15) and others(16) have recognized the exquisite sensitivity of high-risk neuroblastoma to indisulam, with complete responses observed in multiple xenografts, patient-derived xenografts (PDX) and transgenic mouse models. Notably, cells with MYCN amplification exhibit increased sensitivity to aryl sulfonamides, potentially due to their enhanced dependence on RNA processing machinery driven by elevated N-Myc/c-Myc activity, rendering them more susceptible to RNA splicing interference (17). Through the induction of splicing errors in genes within key oncogenic pathways, indisulam can render cancer cells susceptible to additional disruptions caused by other anticancer drugs. For example, we previously showed that several DNA damage repair (DDR) genes are mis-spliced following RBM39 depletion, increasing sensitivity to PARP inhibitors in ovarian high-grade serous carcinoma(18).

In this study, we sought to further define combinatorial vulnerabilities arising from spliceosome disruption through drug-drug library screens combining indisulam with FDA approved anti-cancer agents across various cancer models. This approach revealed a synergistic interaction between indisulam and ALK-inhibitor alectinib, likely due to activity against a recently discovered secondary target of alectinib, RNA Serine-Arginine Protein Kinase 1 (SRPK1) (19). SRPK1 promotes the nuclear translocation and activation of a family of SR splicing factors (SRSFs). The combination of indisulam and alectinib simultaneously depletes RBM39 and inhibits SRPK1-mediated SRSF phosphorylation, resulting in exacerbated RNA splicing defects. The subsequent failure to resolve accumulated DNA damage and R loops results in cell cycle arrest, cell growth inhibition and apoptosis of neuroblastoma cells *in-vitro* and tumour regression *in vivo*. Our study provides important proof-of-concept evidence for the co-targeting of splicing factors as a viable strategy to treat aggressive malignancies such as MYCN-driven high-risk neuroblastoma.

## RESULTS

### Combination drug screens revealed synergy between indisulam and alectinib

We previously demonstrated that indisulam-mediated RBM39 degradation causes wide-spread RNA splicing defects, and an enrichment in aberrant expression of proteins that mediate DNA damage repair and cell cycle regulation, across various cancer types(15,18,20). This led us to speculate that indisulam is likely to generate drug-induced-vulnerabilities that potentiate the efficacy of existing, clinically approved anti-cancer compounds, such as PARP inhibitors(18). RBM39 degraders, such as indisulam and E7820, have been trialed in 47 clinical studies(3), highlighting potential therapeutic combinations ready for clinical evaluation.

To explore this concept, we conducted two drug-library screens combining indisulam with 147 FDA-approved anticancer drugs (NCI plate number #4891, #4892) in an indisulam-sensitive cell line (HCT116, colorectal cancer) and an indisulam-resistant line (KURAMOCHI, ovarian cancer). Drugs were tested at 5 doses (16 nM to 10 µM) and combined with DMSO or indisulam at fixed doses (GI30=50 nM for HCT116, 10 µM for KURAMOCHI) (**Fig. 1a, Supplementary Fig.1a**). Bliss scores(21) were calculated to identify synergistic interactions with indisulam. Depletion of RBM39 by indisulam is dependent on DCAF15 expression, thus we generated CRISPR-Cas9 knockouts of *DCAF15* to identify RBM39-specific drug-drug interactions (HCT116 *DCAF15* wild-type (*DCAF15*^WT^) clone #1 and #2 and *DCAF15* knock-out clones (*DCAF15*^KO^) clone #1 and #2). Drugs with Bliss score>0 in both HCT116-*DCAF15*^WT^ clones were selected as hits which were further validated in triplicate experiments with selected doses in both HCT116-*DCAF15*^WT^ and *DCAF15*^KO^ clones #1 and #2 and ranked by mean Bliss scores of both cell lines. Most of the synergistic drug-drug interactions were rescued in HCT116-*DCAF15*^KO^ cells suggesting that the synergy is dependent on *DCAF15* and RBM39 degradation (**Fig. 1b left**). In KURAMOCHI, the top 20 hits were ranked by mean Bliss scores across all doses (**Fig. 1b right**). The clinical ALK inhibitor alectinib demonstrated strong synergy in both HCT116 (top hit) and KURAMOCHI (second hit) and was therefore selected for further validation using a wider range of drug concentrations. Strong synergistic growth inhibition was observed across a range of doses of indisulam and alectinib in HCT116 (**Fig. 1c**) and KURAMOCHI (**Fig. 1d**), and the synergy was abrogated in HCT116-*DCAF15*^KO^ cells.

**Figure 1.**
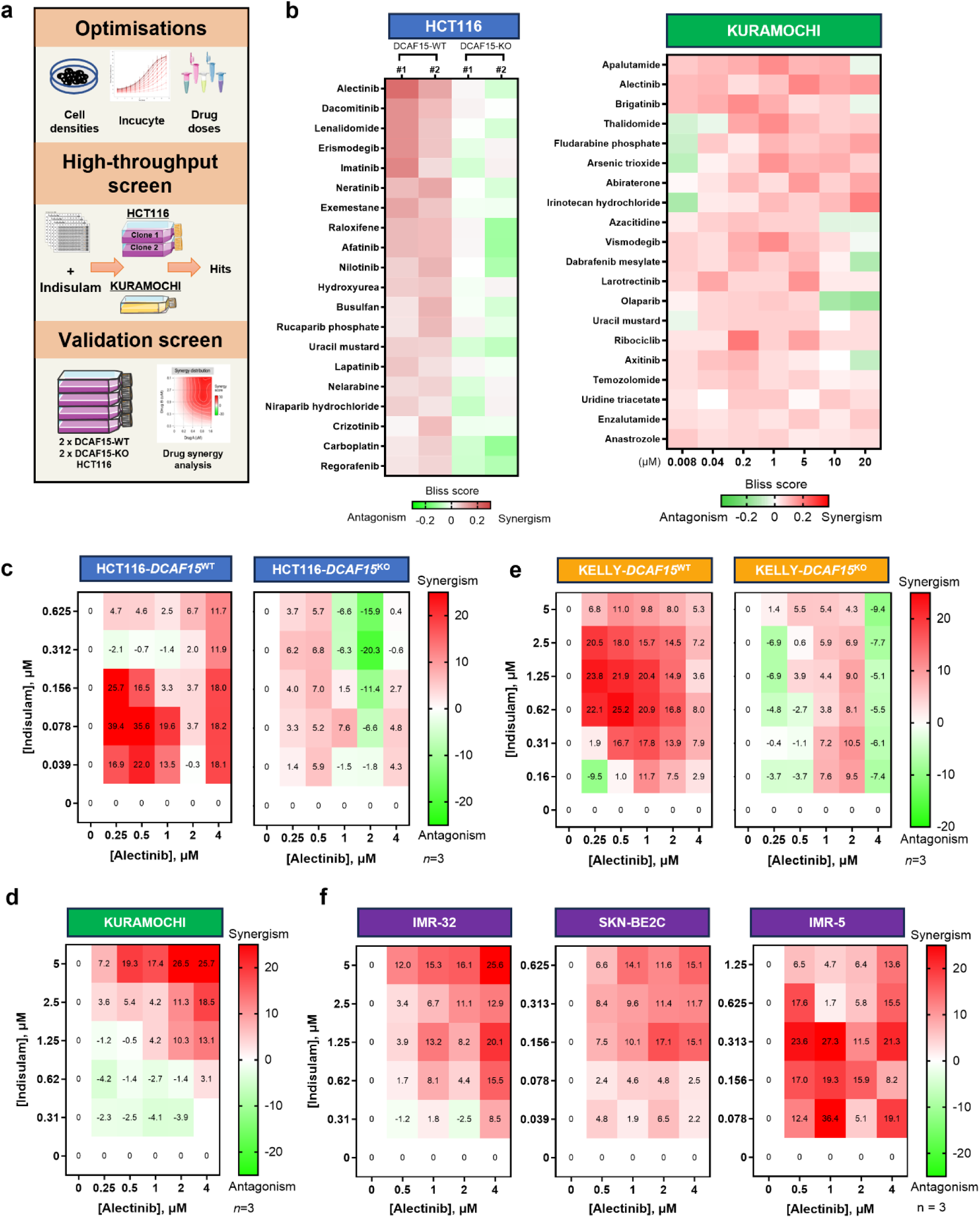
Combination drug screens revealed a synergistic growth inhibition of indisulam and alectinib. a Workflow of combination drug screens in HCT116 and KURAMOCHI using indisulam and 147 FDA-approved anticancer compounds. b Top 20 hits from drug screens in HCT116 (left) and KURAMOCHI (right). HCT116-*DCAF15*^WT^ or HCT116-*DCAF15*^KO^ were treated with selected hits ± indisulam. Bliss scores were calculated from SRB assay data; >0 (red) indicates synergy. KURAMOCHI cells were treated similarly. n=3.; Top 20 hits are ranked by mean bliss score. n=3 replicates. **c-f** Cell lines used for synergy screen (**c, d**) and indicated HRNB cell lines (**e, f**) were treated with alectinib combining indisulam for 72 hours and analysed for cell growth by SRB. Synergy scores were calculated by Synergyfinder (https://synergyfinder.fimm.fi/) using the Loewe method.

These data confirm that combining alectinib with RBM39 degraders yields synergistic growth inhibition, supporting a potential combination therapy across diverse cancer types.

### Indisulam and alectinib exhibit synergistic anticancer efficacy in high-risk neuroblastoma

Considering the potency demonstrated by both indisulam and alectinib as monotherapies in high-risk neuroblastoma(11,15), we further investigated the combination of indisulam and alectinib in four high-risk neuroblastoma cell lines (KELLY, IMR32, SKNBE2C, IMR5) (**Fig. 1e-f**). We demonstrate that combining alectinib with indisulam significantly inhibits cell proliferation compared to monotherapies and display great synergy across a wide-range doses of both compounds on neuroblastoma cell lines, but only additive or antagonistic effects were found in KELLY-*DCAF15*^KO^ cell line confirming the necessity of RBM39 degradation for this pharmacological interaction (**Fig. 1e right**). Similar synergy profiles were observed between alectinib and E7820, a second aryl sulfonamide RBM39 degrader (2)(**Supplementary Fig.1b-c**).

In addition, alectinib treatment alone did not induce cell apoptosis, but significantly increased the percentage of apoptotic cells when combined with indisulam in KELLY and IMR32 (**Fig. 2a-b, Supplementary Fig.2d**). Correspondingly, a significant increase of cleaved caspase 3 was detected in the combination treatment group in KELLY cells (**Fig. 2c**). To evaluate cell cycle progression, we used flow cytometry analysis and detected a notable accumulation of cells in the G2/M phases following 48-hour treatment of indisulam which is consistent with a previous study(16). Co-treatment with indisulam and alectinib increased the percentage of cells in the G1-phase, implying enhanced cell cycle arrest in both G2/M and G1 phase, with a substantial decrease in the percentage of cells in S-phase (**Fig. 2d-e**). These data confirm that indisulam and alectinib act synergistically to suppress the proliferation of neuroblastoma cells, disrupt cell cycle and induce cell apoptosis.

**Figure 2.**
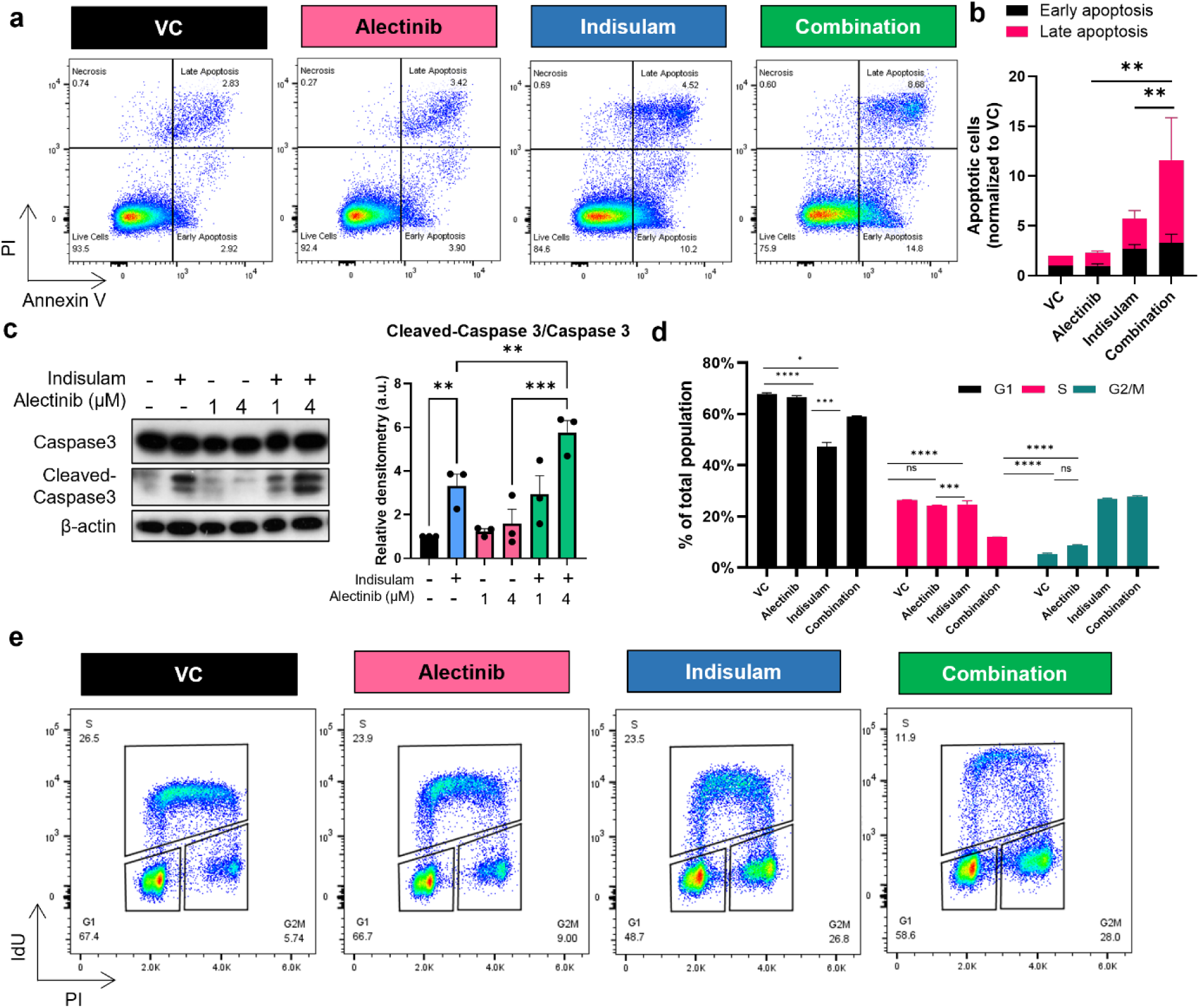
Indisulam and alectinib induce apoptosis and cell cycle disruption in high-risk neuroblastoma. **a-b** Apoptosis analysis via flow cytometry in KELLY cells after 48h treatment with 1.25µM indisulam, 4µM alectinib, or their combination. Cells were labelled by Annexin V and propidium iodide (PI). The significance of difference of total apoptotic cells was calculated by one-way ANOVA. **c** Western blot of KELLY cells treated with indicated doses of indisulam and alectinib for 48 hours. (n=3 independent experiments) significance of difference was calculated by one-way ANOVA. **d-e** Cell cycle analysis by flow cytometry after 48 h treatment with indisulam, Alectinib, or the combination. Cellular DNA content was labelled with PI, replicating cells were labelled with nucleotide analogues 5-Iodo-2-deoxyuridine (IdU). n=3 independent experiments. Two-way ANOVA were employed to determine the statistical significance among the groups. n.s. not significant, * p < 0.05, **p<0.01, ***p < 0.001, ****p < 0.0001.

### ALK inhibition by alectinib is not required for synergistic interaction with indisulam

According to data from the Human Protein Atlas (proteinatlas.org) (22) ALK and RET, both primary targets of alectinib, are expressed in NB cells but not in HCT116 and KURAMOCHI (**Fig. 3a, Supplementary Table 1**) in which synergy was initially discovered. Western blotting confirmed an absence of ALK expression in these cells (**Fig. 3b**). To further test whether the synergy is driven by ALK inhibition, we transfected KELLY, HCT116, and KURAMOCHI cells with an siRNA targeting ALK (siALK) prior to indisulam dosing (**Fig. 3b**). We found no significant difference in cell growth inhibition between siALK and non-targeting control (siNTC) transfected cells (**Fig. 3c**), indicating that ALK inhibition alone does not increase sensitivity to indisulam. We also tested a second ALK inhibitor with a distinct chemical scaffold and binding site, lorlatinib, and found that this compound did not enhance sensitivity to indisulam, except in KELLY cells, which express a mutant, oncogenic form of ALK (ALK^F1174L^) (**Fig. 3d and Supplementary Fig.2a**). However, in cell lines with wild type ALK (IMR32) or lacking ALK expression (HCT116 and KURAMOCHI) there was no sensitization (**Fig. 3d**), consistent with other studies (23), indicating that the efficacy of lorlatinib is specific to ALK mutated neuroblastoma. Further experimental validation is needed to clarify the interaction between inhibition of mutant ALK-driven tumours and RBM39 depletion by indisulam. Nonetheless, these findings suggest that ALK inhibition is not a significant contributor to the synergy we observe between alectinib and the RBM39 degraders indisulam and E7820. This suggests that synergy with RBM39 degradation relates primarily to inhibition of a secondary target of alectinib.

**Figure 3.**
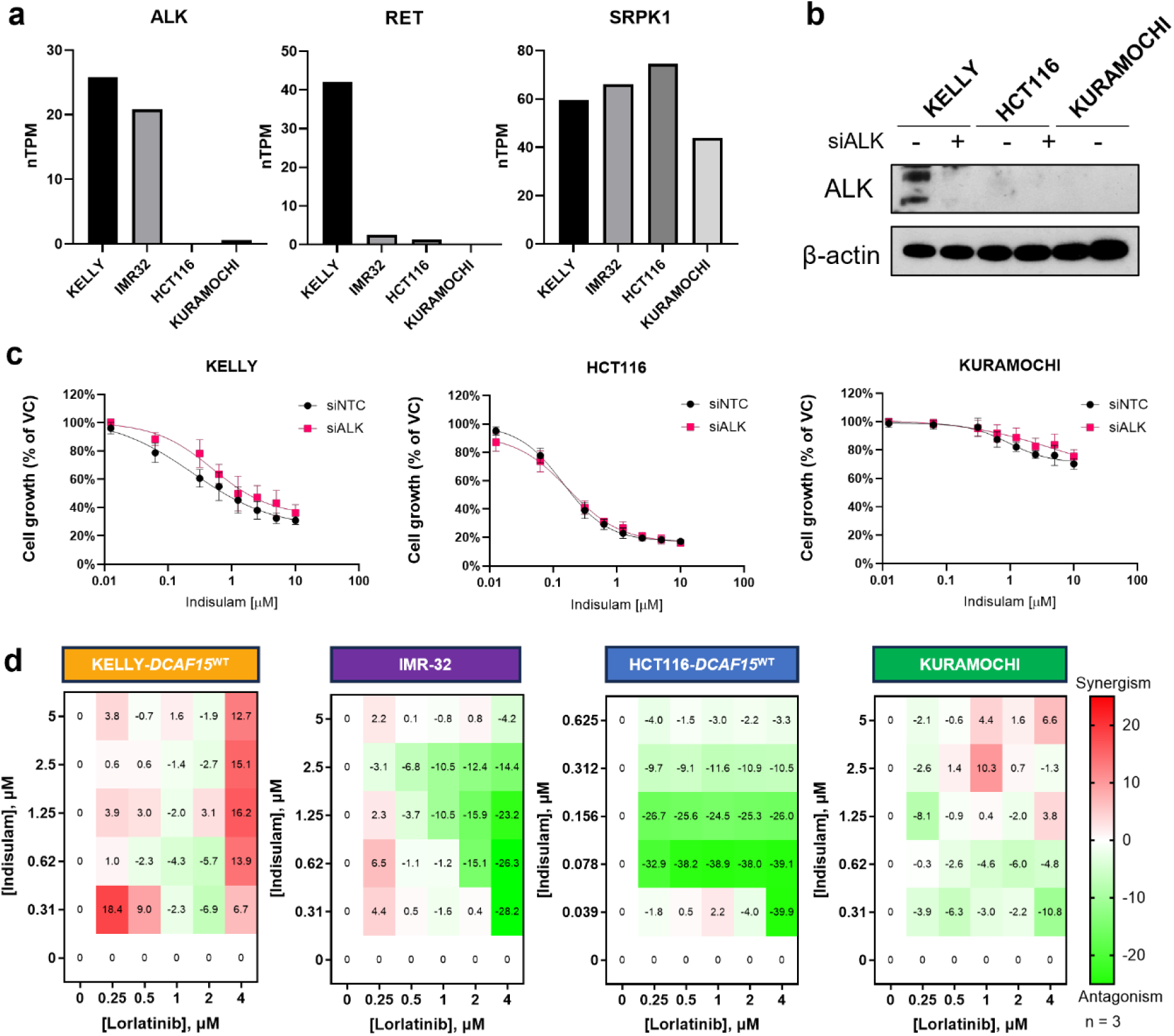
ALK inhibition by alectinib is not an essential determinant of indisulam synergy. **a** RNA expression data as normalized transcript per million (nTPM) values of cancer cell lines were obtained from proteinatlas.org. **b** Western blot confirming the knock-down of ALK gene expression in cells post transfection with siNTC or siALK. **c** Indicated cell lines were transfected with siALK or siNTC for 16 hours, followed by exposure to indisulam for 72 hours. Cell growth was measured by SRB assay. **d** Indicated cell lines were treated with lorlatinib and indisulam concentration matrix for 72 hours. Loewe synergy scores are plotted. n=3 replicates.

### Alectinib exerts potent inhibition of SRPK1

Comprehensive kinase inhibitor screens indicate that alectinib inhibits an array of additional kinases other than ALK (19). The observation that alectinib is a potent inhibition of Serine/arginine (SR) Protein Kinase 1/2 (SRPK1/2) (19,24) is particularly relevant to our research of aberrant RNA splicing. Both SRPK1 and 2 play a central role in regulation of alternative splicing via phosphorylation and activation and nuclear translocation of the Serine/Arginine-rich Splicing Factors (SRSFs) (25,26). In MYCN-amplified neuroblastoma, SRPK1 expression is elevated and associated with reduced survival rates (**Supplementary Fig.2b-c**). In contrast, lower survival rates are not associated with increased expression of SRPK2 nor SRPK3, indicating an increased dependency on SRPK1 to drive tumour growth in MYCN-amplified tumours. Furthermore, a genome-wide CRISPR study showed that loss of SRPK1 sensitizes to indisulam treatment in various solid tumour cell lines (27). Thus, we further explored the treatment combination of SRPK1i and RBM39 degraders. To this end, we assessed the synergy between indisulam and SPHINX31, a known SRPK1 inhibitor(28). Co-treatment of SPHINX31 and indisulam exhibited synergistic growth inhibition cancer across all three cell lines (**Fig. 4a**). Time-lapse cell confluency monitoring in KELLY-*DCAF15*^WT^ and *DCAF15*^KO^ showed that co-treatment significantly suppressed the growth of KELLY-*DCAF15*^WT^, compared to individual drug treatments. Conversely, in the absence of *DCAF15*, the combination therapy displayed similar potency to SPHINX31 monotherapy (**Fig. 4a**), suggesting this interaction is DCAF15-dependent and thus RBM39-dependent.

**Figure 4.**
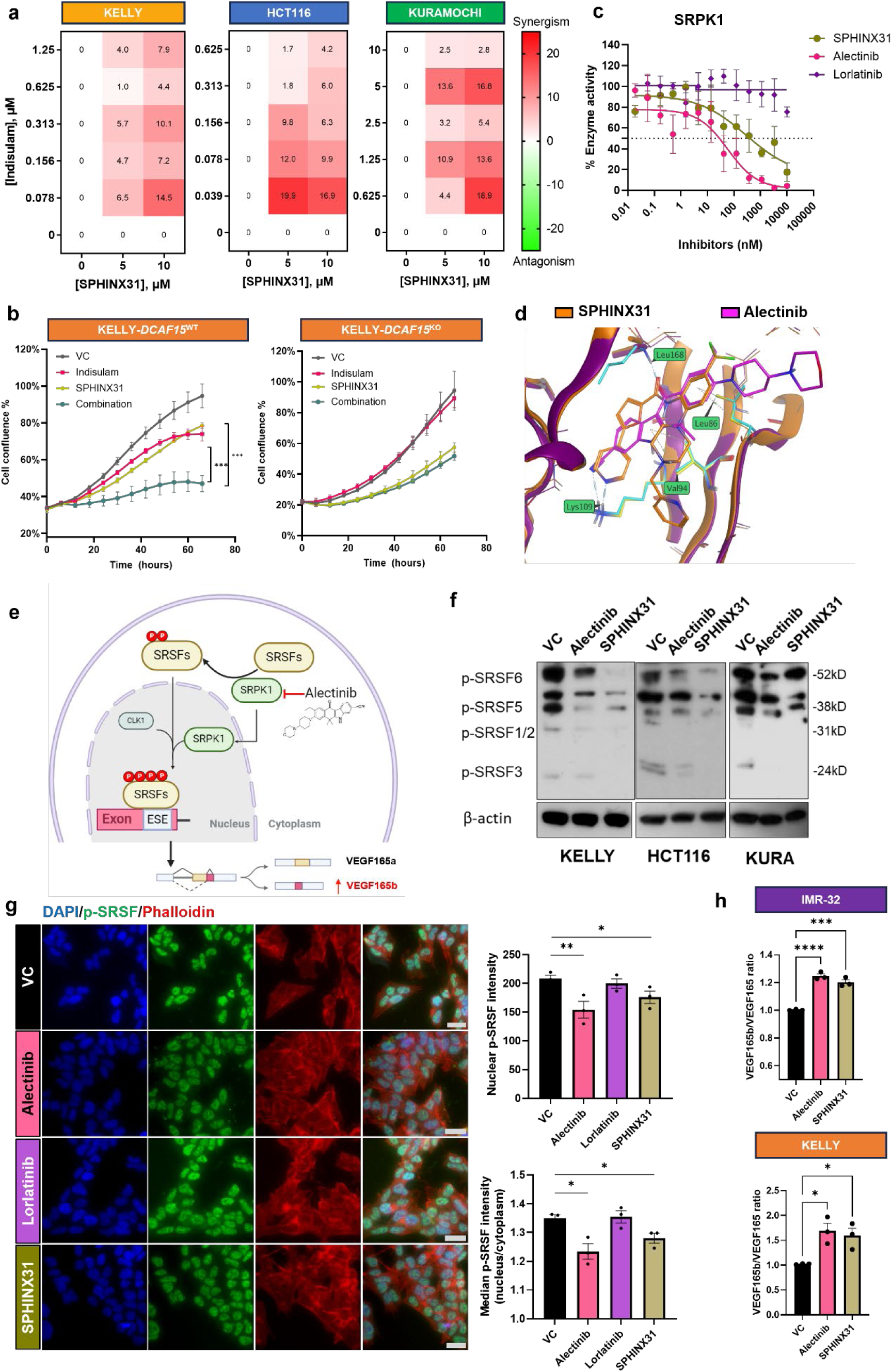
SRPK1 inhibition is synergistic with indisulam in high-risk neuroblastoma. **a** Indicated cell lines were treated with indisulam and SPHINX31. Loewe synergy scores as indicated. **b** KELLY-*DCAF15*^WT^ and *DCAF15*^KO^ cells were treated with indisulam and SPHINX31 individually or in combination for 72 hours. Cell confluence was monitored every 6 hours using the Incucyte live cell imaging system. One-way ANOVA was used to determine significance. **c** SRPK1 enzyme activity measured with ADP-Glo™ Kinase Assay with indicated doses of compounds. **d** Alectinib (pink) docked into SRPK1 complexed with SPHINX31(orange). **e** Diagram depicting Alectinib inhibition against SRPK1. Made with Biorender. **f** Western blotting of p-SRSF in KELLY, HCT116 and KURAMOCHI post-treatment. **g** Immunofluorescent staining of p-SRSF in KELLY post-treatment. Nuclear intensity and cytoplasmic staining intensity are calculated using DAPI and Phalloidin (F-actin) segmentation. Scale bar = 20μm. **h** qRT-PCR of *VEGF165b* and total *VEGF165* in KELLY and IMR32 cells normalised to GAPDH. * p < 0.05, **p < 0.01, ***p < 0.001. n = 3 independent experiments.

The inhibition of SRPK1 by alectinib was confirmed by enzyme activity assays which reported a half-maximal inhibitory concentration (IC50) of 26nM. This inhibition is more potent than that of SPHINX31 (IC50 = 375nM) with lorlatinib showing very little inhibitory action (**Fig. 4c**). Comparison of the crystal structures of the SRPK1-Alectinib and SRPK1-SPHINX31 complexes confirmed that alectinib and SPHINX31 interact with the ATP-binding groove of SRPK1, with reasonable affinity and occupying similar positions within the binding site (**Fig. 4d** and **Supplementary** Fig.3). Notably, alectinib inhibited SRPK2 with an IC50 of 700nM while SPHINX31 showed no inhibitory effect (**Supplementary Fig.2d**).

SRPK1 phosphorylates SRSFs, triggering their nuclear translocation and activity in modulating RNA splicing (**Fig. 4e**). Both alectinib and SPHINX31 reduced phosphorylated SRSF (p-SRSF) in KELLY, HCT116 and KURAMOCHI cells (**Fig. 4f)** and decreased the nuclear-to-cytoplasmic ratio of p-SRSF intensity within 4 hours in KELLY cells, while lorlatinib lacked this effect (**Fig. 4g**). SRSFs are known to promote selection of the proximal splice site in *VEGF* pre-mRNA and inhibiting SRPK1 alters *VEGF* splicing to generate the *VEGF165b* isoform(19,29). Treatment of both alectinib and SPHINX31 resulted in a switch and increased *VEGF165b* mRNA in high-risk neuroblastoma cells (**Fig. 4h**).

In conclusion, our findings reveal that alectinib is a potent inhibitor of SRPK1, consequently modulating downstream function and RNA splicing through a reduction in the phosphorylation of SRSF splicing factors. Notably, alectinib demonstrates more robust inhibition of SRPK1 as compared to the selective inhibitor SPHINX31, an intriguing observation that could potentially expedite the clinical repurposing of alectinib as an SRPK1 inhibitor.

### Alectinib and indisulam exacerbate RNA splicing defects, increase R-loop formation and DNA damage

Proteomic data from our previous study (15) showed that indisulam exposure leads to an increase in protein expression of most SRSF family members (**Supplementary Fig.4a**). Here, we observed enhanced nuclear translocation of p-SRSF post-indisulam treatment, which was rescued in *DCAF15*^KO^ cells (**Supplementary Fig.4b-c**), suggesting that SRPK1/SRSF pathway activation is a compensatory mechanism following RBM39 degradation. RBM39 has previously been identified as a potential SRPK1 target [25] and we observed a direct interaction between RBM39 and SRPK1 in neuroblastoma cells via Proximity Ligation Assay (PLA) (**Supplementary Fig.4d**). This supports our hypothesis that the synergy between alectinib and indisulam arises from their combined disruption of RNA splicing, by dually targeting interdependent spliceosomal components, thereby limiting cellular adaptation.

To evaluate the impact of treatment on alternative splicing, IMR32 cells were treated with alectinib, indisulam, or both for 24 hours. RNA-seq and rMATS analysis revealed that while indisulam induced widespread splicing changes, particularly skipped exons (SE) and that alectinib alone had a milder effect, combination treatment resulted in additional 7,646 altered events (33%) and 1,758 mis-spliced genes (25%) beyond an additive effect (**Fig. 5a)**. Notably, alectinib-affected genes largely overlapped with indisulam targets **(Fig. 5b)**, further supporting the likelihood of functional cooperation between RBM39 and SRPK1. Gene ontology (GO) analysis of alternatively spliced genes revealed enrichment in pathways regulating genome stability, including *DNA repair*, *chromatin remodeling* and *DNA damage response* (**Fig. 5c)**. The significantly affected AS events were plotted following combination treatment. (**Fig. 5d)**. Several most affected events were validated through custom PCR primer design of affected exon regions (**Fig. 5e-f)**, confirming that splicing disruption was significantly enhanced when combining RBM39 degraders and SRPK1 inhibition in both KELLY and IMR32 cells (**Fig. 5e-f, Supplementary Fig.5a**); such an effect was not observed with lorlatinib or siALK (**Supplementary Fig.5b-d**). These results suggest the synergy observed between alectinib and indisulam may be attributed to the compound reduction of canonical isoform expression and enhanced functional loss in effected pathways, such as DNA damage repair. For example, expression of the canonical CHK1 isoform was nearly abolished by the co-treatment of indisulam with either alectinib or SPHINX31. Moreover, CHK1 protein was depleted following indisulam and alectinib combination treatment (**Supplementary Fig.5e**),

**Figure 5.**
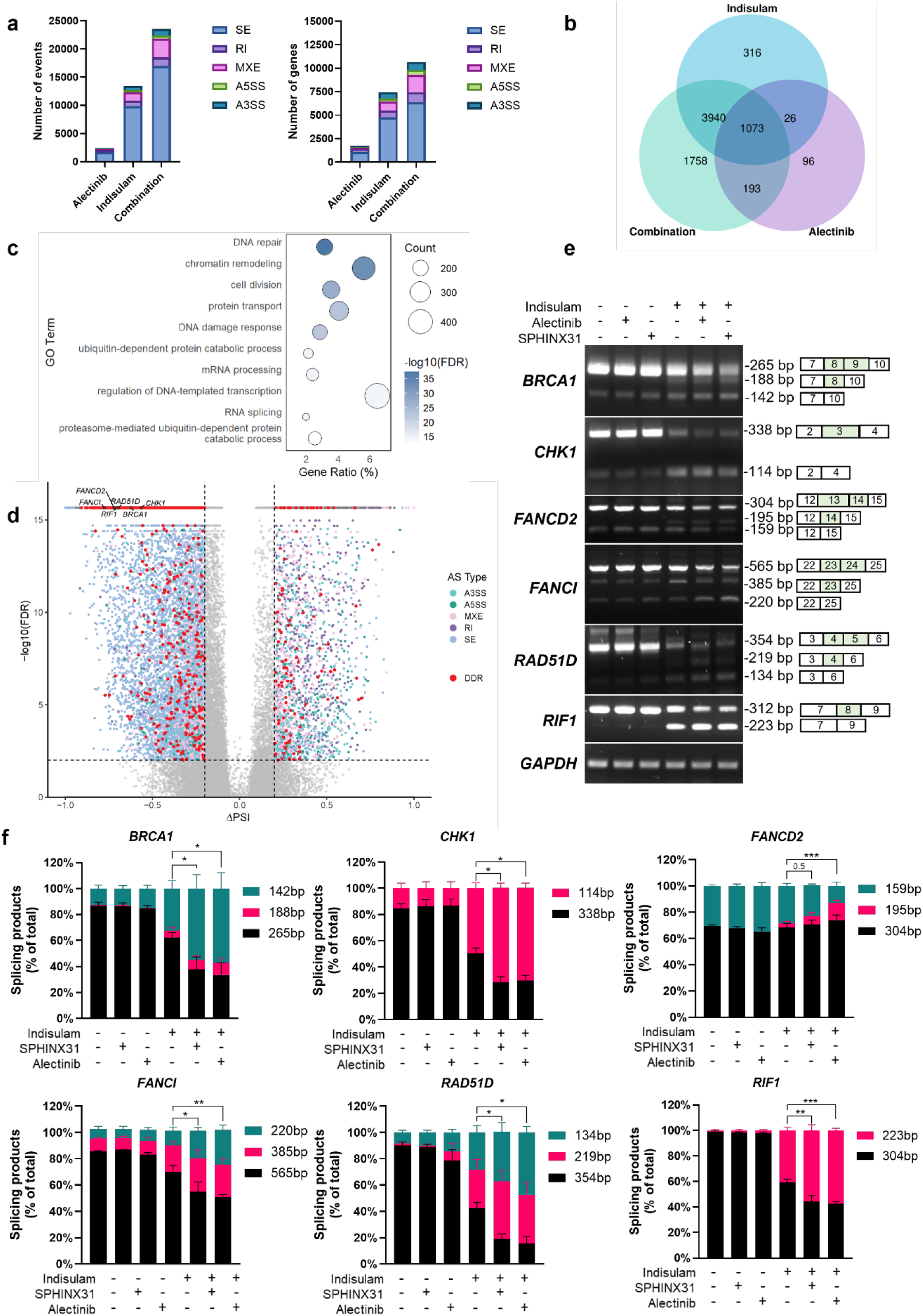
Alternative-splicing elicited in neuroblastoma cells by treatment with alectinib, indisulam and the combination. **a** Number of alternative splicing (AS) events (left) and affected genes (right) detected by rMATS, classified into exon skipping (SE), retained introns (RI), mutually exclusive exons (MXE), alternative 5′ splice sites (A5SS), and alternative 3′ splice sites (A3SS). **b** Venn diagram showing overlap of alternatively spliced genes across treatments. **c** Gene Ontology (GO) enrichment analysis of AS genes in the combination group; dot size reflects gene count, colour indicates –logFDR. **d** Volcano plots display –log₁₀(FDR) values calculated by rMATS. Events with FDR = 0, due to numerical underflow from highly significant p-values, were assigned a nominal value of 1e–16 for visualisation purposes. Significant splicing events were defined as those with FDR < 0.01 and |ΔPSI| ≥ 0.2. DDR related genes are highlighted in red, and six of these genes were selected for further experimental validation (labelled). **e** PCR validation of exon skipping events in DDR genes (representative image of n=3). *GAPDH* was included as a control. **f** Densitometry analysis of alternative splicing products, Two-way ANOVA was used to detect the significance.

To determine whether splicing-induced depletion of DDR genes increased DNA damage, we assessed γH2AX levels as a marker for DNA double stranded breaks(30). Immunofluorescence and western blot showed significant γH2AX accumulation following combination treatment compared to monotherapies (**Fig. 6a-b, Supplementary Fig.6a**). Importantly, this increase was absent when indisulam was combined with lorlatinib (**Supplementary Fig.6b-c**), indicating that the effect stems from RNA splicing disruption rather than ALK inhibition.

**Figure 6.**
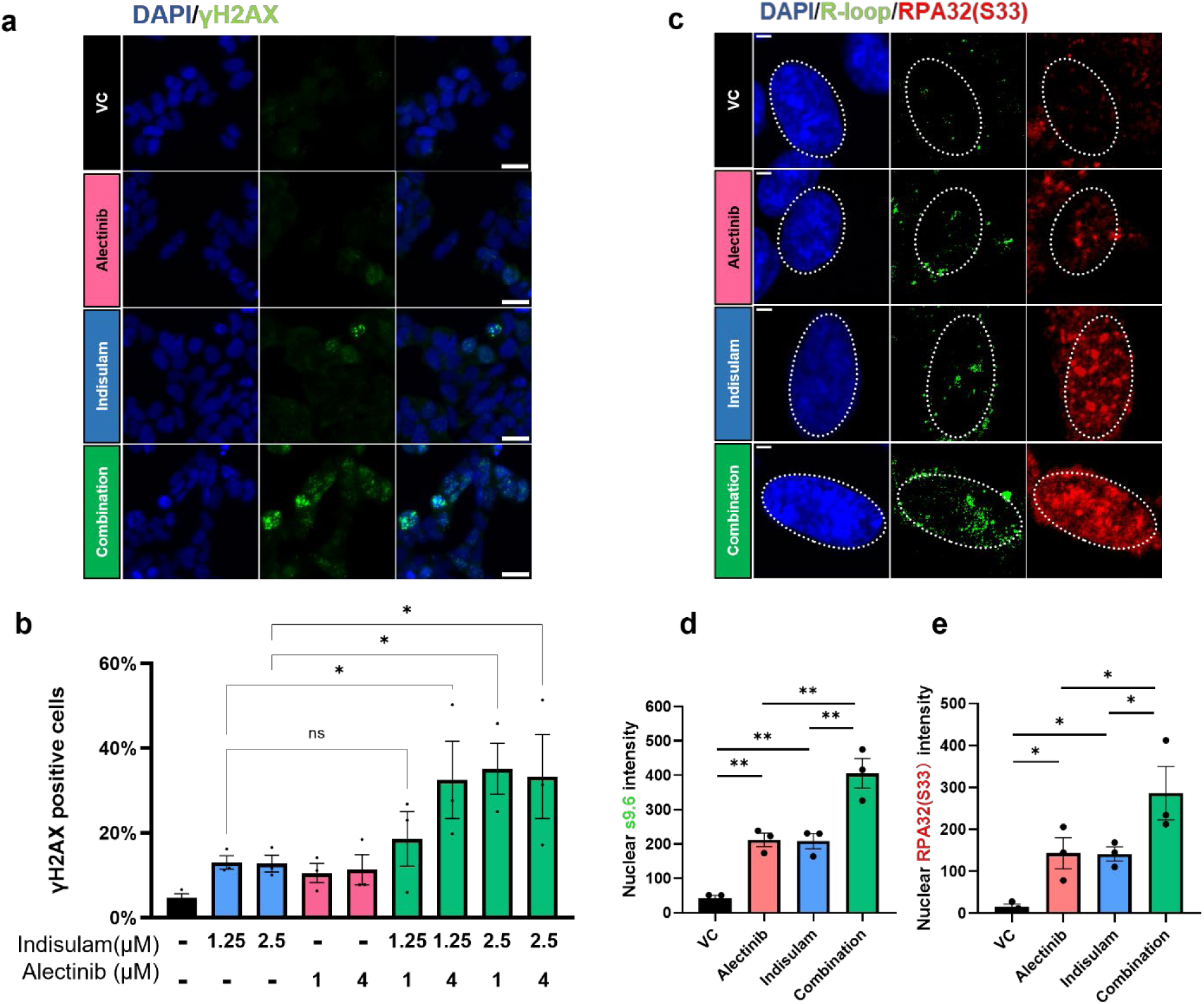
Combination of indisulam and alectinib induce DNA damage, R-loop accumulation and replication stress. **A-b** Immunofluorescence staining of γH2AX in KELLY cells treated with indisulam in combination of alectinib. Representative images of KELLY treated with 4μM alectinib and 1.25μM indisulam are shown. Scale bars = 20μm. Nuclear foci >5 indicates positive cells. n=3 independent experiments. **c** Representative images for nuclear S9.6 and RPA32 (S33) foci in KELLY cells, n = 3 independent experiments, Scale bars = 2.5μm. **d-e** Quantification of nuclear S9.6 intensity d and pRPA32 e intensity in KELLY cells. Cells were treated with 4μM alectinib monotherapy or with 1.25μM indisulam; nuclear signal intensity were measured by merging to DAPI; n>300 cells from 3 independent experiments were quantified. Significance was analysed among the median values.

R-loops are DNA:RNA hybrids that are a product of intermediate transcription and must be resolved to permit progression of replication forks (31,32). R-loop accumulation results in transcription-replication collisions, which generate replication stress and double-strand breaks (33). Many R-loop regulatory genes were mis-spliced after combination treatment. For example, BRCA1 interacts with the putative RNA helicase Senataxin (SETX), forming a complex that localizes to R-loops(34) and Fanconi Anemia genes recruit RNA processing enzymes to resolve R-loops generated by moderate replication stress(35), while RIF1 facilitates replication fork protection and efficient restart to maintain genome stability(36). Notably, SR proteins regulated by SRPK1 can also impede R-loop formation(31). Dysregulation of these functions lead to accumulation of unresolved R-loops, replication fork breakage, and genome instability (31).

We quantitated R-loops using S9.6 antibody(37). Treatment with either alectinib or SPHINX31 led to increased nuclear S9.6 staining indicating the accumulation of R-loops. The combination treatment significantly increased R-loop formation as compared to monotherapies (**Fig. 6c-d, Supplementary Fig.6d-f**). To evaluate replication stress induced by Rloop accumulation(31,38), we quantified the intensity of phosphorylated RPA32 serine 33 (pRPA), a marker of single-stranded DNA (ssDNA) at stalled replication forks. We observed an increase following exposure to the combination treatment relative as compared to monotherapy (**Fig. 6e, Supplementary Fig.6e**). DNA fibre analysis using BrdU and IdU labelling further revealed a reduction in track length and the significantly decreased IdU/BrdU ratio following combination treatment, indicating increased fork stalling and replication stress. Collectively, these findings suggest that dual targeting of SRPK1 and RBM39 leads to the accumulation of unresolved R-loops, promoting replication stress, and contributing to subsequent DNA damage in high-risk neuroblastoma.

### Combination of indisulam and alectinib improves survival in a Th-MYCN/ALK^F1174L^ neuroblastoma model

Gain-of-function mutations in ALK account for 10-15% of high-risk neuroblastoma (6). ALK inhibitors have been shown to be safe and effective in treating ALK-driven neuroblastoma in a Phase I trial(39,40), and are currently being investigated in several Phase III trials(39,41,42), highlighting their potential for clinical use in ALK-positive neuroblastoma. Considering the clinic relevance and benefit, we used an allograft mouse model implanted with Th-MYCN/ALK^F1174L^ tumour cells derived from genetically engineered mouse models (GEMMs) (**Fig. 7a**).

**Figure 7.**
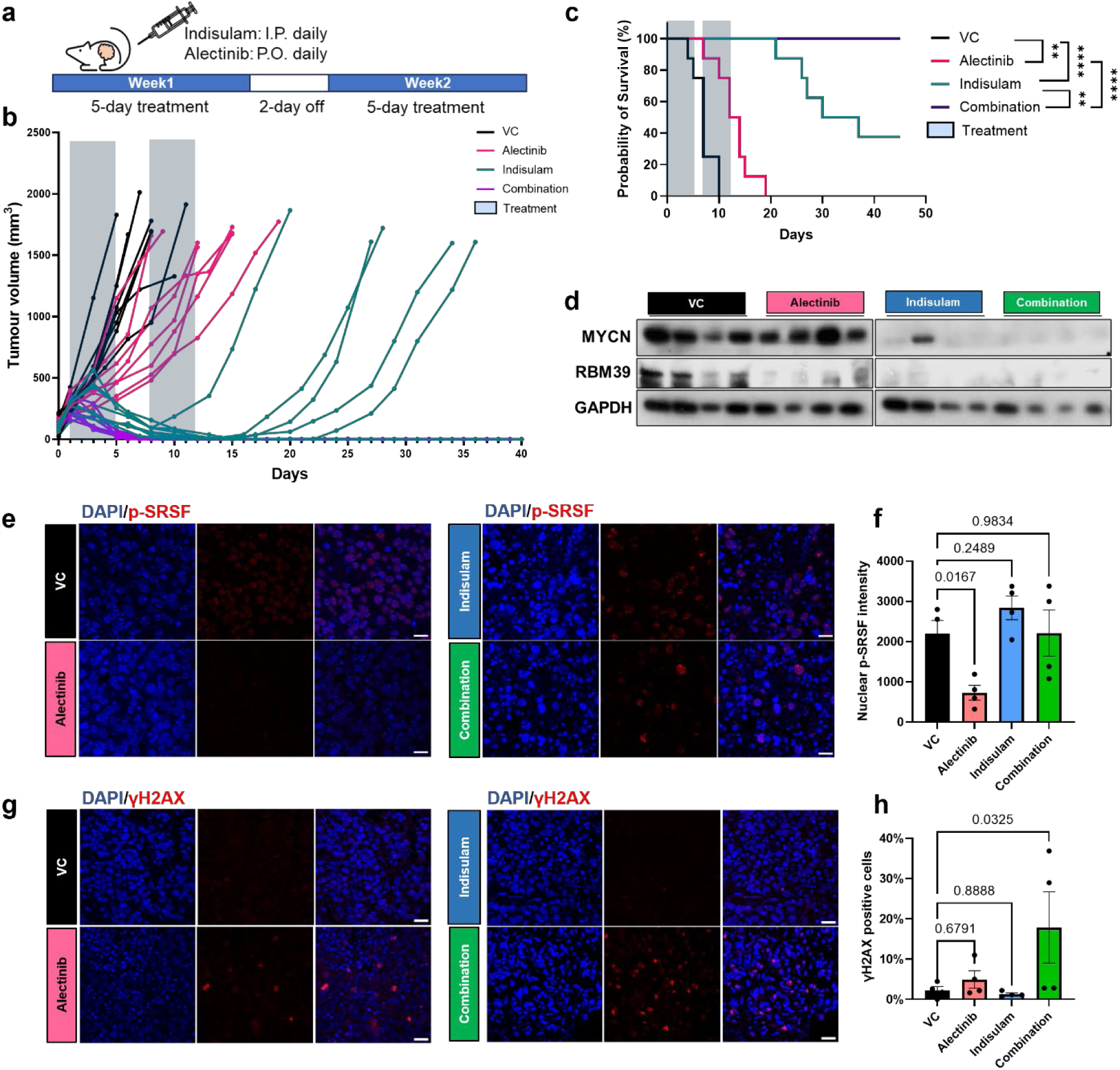
Combination of indisulam and alectinib suppress neuroblastoma tumour growth. **a** Th-ALK^F1174L^/Th-MYCN GEMM tumour cells were injected into a cohort of mice subcutaneously, intravenous (i.v.) administration of 10 mg/kg of indisulam and oral gavage (p.o.) 40 mg/kg of alectinib for 2 weeks with 5 days on and 2 days off. **b** Tumour volume measured 3 times a week until the humane end-point was reached. Cyan shade indicates the treatment period. n = 8 for each group. **c** Survival plot of mice bearing Th-ALK^F1174L^/Th-MYCN allografts treated with vehicle or alectinib or indisulam or combination therapy. **d** Western blotting in tumour tissues after administration of vehicle or alectinib or indisulam or combination therapy for 3 days (n = 4 per group). **e-h** Immunofluorescence staining was performed on tumour sections, with DAPI used for nuclear staining. Nuclear p-SRSF intensity (E-F) was measured by merging to DAPI. Nuclear γH2AX (G-H) foci >5 indicates positive cells. Scale bar = 20 µm. n.s. not significant, * p < 0.05, **p<0.01, ***p < 0.001, ****p < 0.0001.

Tumour-bearing mice were treated (indisulam 10mg/kg, alectinib 40mg/kg or combination) for 2-weeks with a 45-day follow-up. Both monotherapies significantly enhanced survival however all alectinib treated mice relapsed on treatment and 5/8 mice in the indisulam monotherapy group relapsed after withdrawal of treatment. Complete tumour regression without subsequent relapses resulting in a 100% survival rate were reported in the combination group (**Fig. 7b-c**) consistent with the increased efficacy in-vitro.

Tumour samples were collected for pharmacodynamic analysis after 3 days of treatment. Western blotting showed a clear depletion of RBM39 following indisulam and combination therapy, confirming the on-target effect of indisulam (**Fig. 7d**). Furthermore, treatment with 10 mg/kg indisulam reduced MYCN expression, as previously reported by Singh et al (43), suggesting a potential link between RBM39 inhibition and suppression of MYCN (**Fig. 7d**). Additionally, a decrease in p-ALK **(Supplementary Fig.7a)**, and a reduction in p-SRSF (**Fig. 7f**) confirmed the on-target effects of alectinib on SRPK1 and ALK. Double strand breaks measured by accumulation of γH2AX foci, were increased in combination therapies (**Fig. 7g**), in line with our observations *in-vitro.* However, indisulam alone did not significantly induce γH2AX accumulation, possibly due to the transient nature of γH2AX focus formation during the DNA damage response. However, clear nuclear fragmentation was observed in mice treated with indisulam and combination therapy (**Fig. 7g**). Importantly, no significant weight loss was observed during the trial, indicating no significant toxicity (**Supplementary Fig.7b)**, and we observed no significant RBM39 loss in other organs after 3-day treatment (**Supplementary Fig.7c**).

Taken together, these data confirm that in neuroblastoma the therapeutic benefit of RBM39-depleting agents can be enhanced through combination with SRPK1 inhibitors, such as alectinib. This study provides proof-of concept for the co-targeting of functionally interdependent splicing factors using repurposed clinical agents to more effectively target aggressive childhood cancers such as neuroblastoma. The strategy could apply more widely to other oncogene-driven cancers that exhibit a high dependency on faithful splicing of mRNA.

## DISCUSSION

Indisulam has been recently identified as a molecular glue capable of regulating RNA alternative splicing(2). By inducing splicing errors of genes in key oncogenic pathways, indisulam and other spliceosomal inhibitors may render cancer cells susceptible to additional disruptions caused by other anticancer drugs. Thus, it holds potential for implementation as combination therapy. We performed extensive drug screens with FDA-approved anticancer agents which revealed several potential drug-drug interactions with indisulam. Notably, the synergy between indisulam and PARP inhibitors, also observed in our screens, has been corroborated by other studies in high-grade ovarian carcinoma(18,20), colorectal cancer, and breast cancer(44), giving confidence in our screening approach. Our study also highlights the fact that such screens can reveal beneficial interactions based on polypharmacology(engagement of targets beyond the primary focus of clinical development)(45), presenting novel rationales for drug repurposing.

High-risk neuroblastoma, which constitutes about 15% of all neuroblastoma cases, is characterized by aggressive behaviour and poor prognosis(7). Gain-of-function mutations in ALK account for 10-15% of primary tumours (6), hence a number of trials have investigated the use of alectinib and other ALK inhibitors in this clinical setting. While the more selective ALK inhibitor loratinib has shown greater traction for the treatment of mutant ALK-driven neuroblastoma(39), treatment resistance remains a significant clinical challenge(46). Our study demonstrates that the activity of alectinib against SRPK1 can be exploited therapeutically in combination with other spliceosomal inhibitors. This presents an interesting strategy to treat patients with emerging resistance to ALK inhibitor therapy, including in lung cancer where alectinib has demonstrated much greater clinical use(41) and beyond into other cohorts irrespective of ALK status. While other inhibitors of SRPKs and related SRSF regulators such as Cirtuvivint(47) and TG-003(48) are currently in early development, the fact that an established therapeutic tool can be repurposed in this way could accelerate the clinical testing of SRPK1/2 inhibition and co-targeting of splicing factors as an anti-cancer treatment strategy, particularly in tumours where an ALK inhibitor might normally be indicated.

While we have successfully demonstrated the principle of co-targeting functionally interdependent splicing as an anticancer strategy, several resulting questions arise. Our investigation into the functional consequences of combined RBM39 degradation and SRPK1 inhibition underscores their interdependency in regulating alternative splicing, but it is not yet clear if RBM39 is a direct target of SPRK1 as has been suggested by global interaction profiling (49) and our co-localisation experiments. More broadly, we need further investigation to understand the interactions and co-regulation of different splicing factors that are also cancer targets such as RBM39 and SPRK1, and to develop more detailed mechanistic models of how these regulate alternative splicing together.

Interestingly, RNA-seq analysis revealed dual targeting SRPK1 and RBM39 synergiscally amplifies RNA splicing dysfunction and downstream cellular stress. Targeting functionally connected nodes within the splicing machinery may therefore offer an effective strategy to overcome cellular adaptation and enhance therapeutic efficacy in high-risk neuroblastoma. However, as with the targeting of spliceosomal components in general, it is unclear if the sequelae of accumulating R loops, cell cycle arrest, DNA damage and eventual cell death that we observe in combined SRPK1/RBM39 interference are lineage-dependent, driven by the mis-splicing and functional loss of a select set of critical genes, or the cumulative consequence of widespread alternative splicing events across multiple functional pathways. Further work to resolve such mechanistic questions will be important to understand what the most relevant pharmacodynamic readouts will be for future discovery experiments and which predictive biomarkers can best identify patients most likely to benefit from such combinations (i.e. for stratification).

In summary, the present study supports a model in which co-treatment indisulam and alectinib synergistically impacts RNA splicing through simultaneous RBM39 degradation and SRPK1 inhibition. This dual RNA splicing interference exacerbates splicing errors in key genes regulating genome stability, leading to R-loop accumulation and increased DNA damage. These events eventually result in cellular apoptosis and growth inhibition in high-risk neuroblastoma *in vitro*, as well as suppression of tumour growth *in vivo*. These insights shed new light on translating research on spliceosome inhibition into clinical applications, ultimately aiming to improve the prognosis for patients with high-risk neuroblastoma.

## MATERIALS AND METHODS

### Cell lines and cell culture

Human neuroblastoma cell lines were cultured in MEM (Gibco™) with 10% fetal bovine serum (FBS), 1% penicillin/streptomycin (P/S), 1 mM sodium pyruvate, and 2 mM L-glutamine. KURAMOCHI was maintained in DMEM with 10% FBS, 2 mM L-glutamine, 5.6 mM glucose, and 1% P/S. HCT116 was cultured in RPMI 1640 with 10% FBS, 2 mM L-glutamine, and 1% P/S. All cells were authenticated and routinely tested for mycoplasma. Cells were maintained at 37°C with 5% CO₂.

### Sulforhodamine B (SRB) assay

Cells were fixed with 10% cold trichloroacetic acid, stained with 0.4% SRB for 30 min, and washed with 1% acetic acid. Bound dye was solubilized in 10 mM Tris-base, and OD measured at 565 nm. Results were normalized to vehicle control.

### Synergy analysis

Cell growth inhibition was assessed by SRB assay after 72 hours of drug exposure. The Loewe additivity model was used to describe the interaction between two agents[32]. Synergy scores were calculated using SynergyFinder web application (RRID:SCR_026127) to compute synergy scores and the Loewe model was selected. Scores >10 indicate synergy, −10 to 10 additivity, and <−10 antagonism.

### siRNA Transfection

Cells were seeded in 96-well plates and cultured overnight before transfection. ON-TARGETplus Non-targeting Control Pool (Horizon Discovery, D-001810-10-05) and ON-TARGETplus siALK (Horizon Discovery, L-003103-00-0005) (30 nM final concentration) were delivered using Lipofectamine® 2000 (Invitrogen, 11668-019) in Opti-MEM (Gibco), following the manufacturer’s instructions. Opti-MEM only and Lipofectamine-only conditions were included as controls. Cells were incubated for 16 h prior to further treatment..

### Western blotting

Whole-cell lysates were prepared using RIPA buffer (Sigma Aldrich) with 1% HALT protease and phosphatase inhibitor cocktail (Thermo Scientific). Equal amounts of protein from each sample were separated on a 4-20% Bis-Tris SurePAGE gel (GenScript) and then transferred onto PVDF membranes. The membranes were blocked with 5% milk for 1 hour at room temperature, followed by an overnight incubation at 4°C with CHK1 (Cell Signaling Technology Cat# 2360, RRID:AB_2080320), p-CHK1 (Cell Signaling Technology Cat# 2348, RRID:AB_331212), γH2AX (Sigma-Aldrich Cat# 05-636, RRID:AB_309864). HRP-conjugated secondaries and chemiluminescent substrate were used for detection.

### End-point and quantification PCR reactions

1500ng RNA was utilized for reverse transcription. The resulting cDNA templates were subsequently employed for qPCR using thePowerTrack SYBR Green Master Mix (Applied Biosystems) or End-point PCR utilizing the Q5® High-Fidelity PCR Kit (New England Biolab). PCR products were resolved on 2% agarose gels and quantified with Fiji V2.0.0.

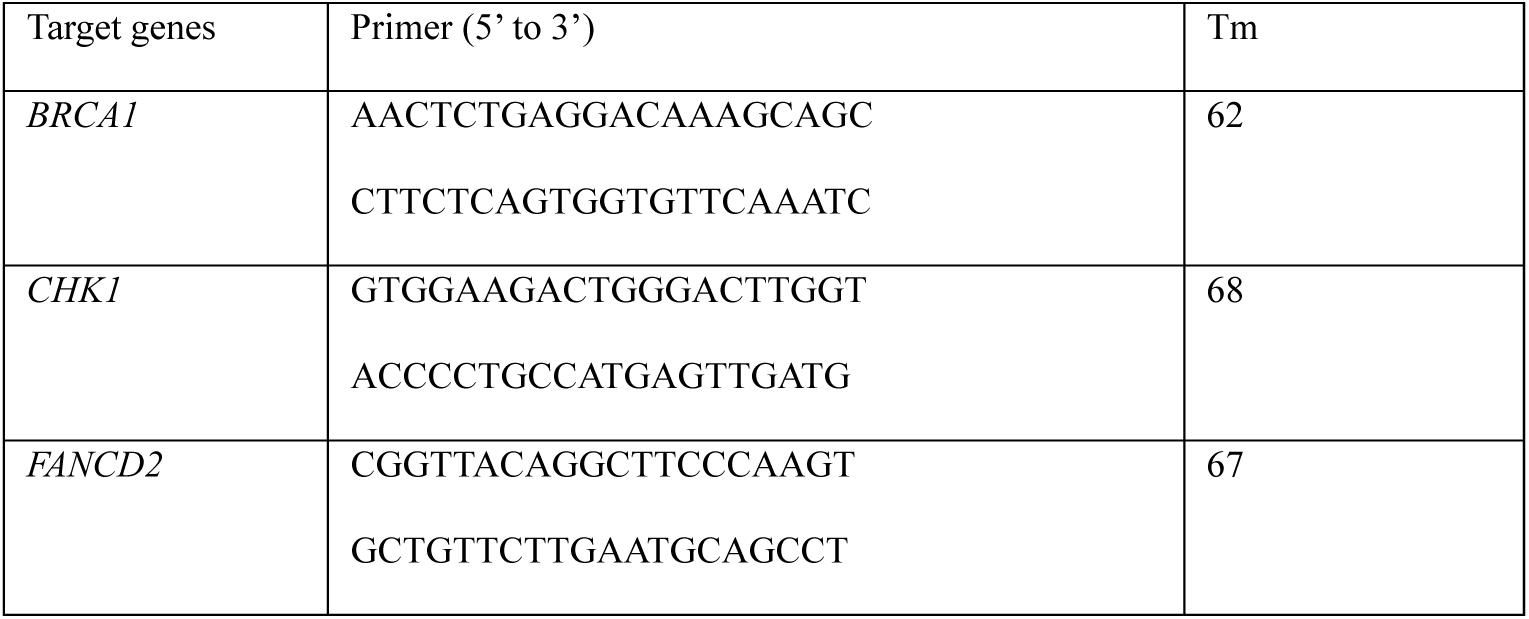
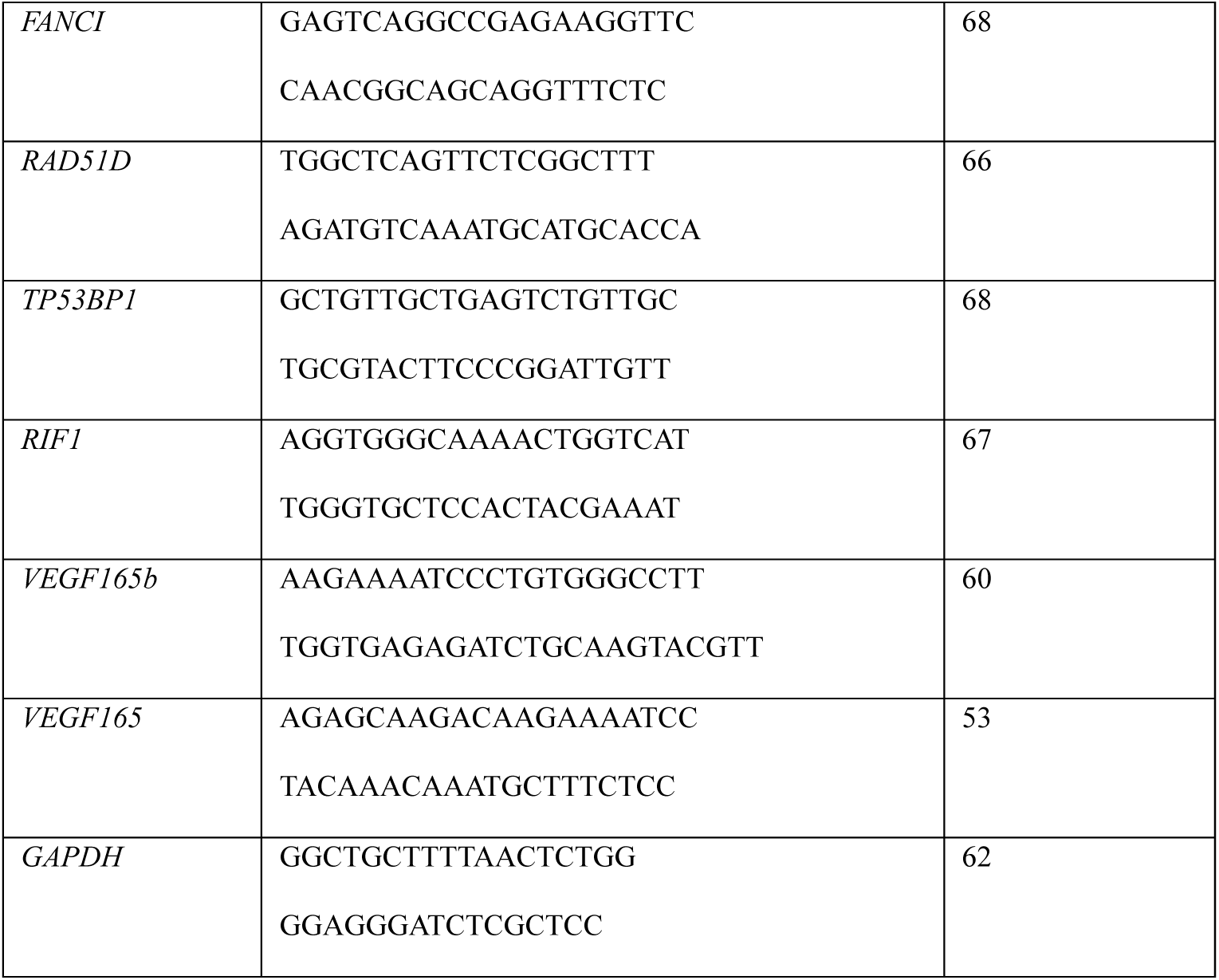

### Immunofluorescence staining

KELLY cells were fixed with 4% PFA, permeabilized with 0.1% Triton-X100, and blocked in 5% BSA. Subsequently, they were incubated overnight at 4°C with anti-phosphoepitope SR proteins antibody (1:100, Millipore Cat# MABE50, RRID:AB_10807429) or γH2AX (1:500, Sigma-Aldrich Cat# 05-636, RRID:AB_309864) in blocking buffer ,and then stained with a mix of Alexa Fluor 488-conjugated goat anti-mouse antibody (1:500, AF488, Thermo Scientific), DAPI (1μg/ml, Biotium, Cat#40043), and Phalloidin-647 (1:50, Cell Signaling Technology Cat# #8940S) for 1 hour at room temperature. The cells were washed again three times with PBS-T before imaging with an INCell Analyzer 1000 microscope (GE Healthcare) at 20x magnification.

S9.6 (1:250, Millipore Cat# MABE1095, RRID:AB_2861387) and phospho RPA32-S33 (1:250, Novus Cat# NB100-544, RRID:AB_10002227) were detected in KELLY cells fixed with cold methanol at -20°C for 10 minutes and slides were imaged using the Aurox laser-free confocal microscope. To confirm the specificity of the S9.6 antibody, RNase H (Ribonuclease H, 1:100, NEB Cat#M0297S) was applied to the fixed cells before antibody incubation.

Image analysis was performed using Cell Profiler V4.2.6, with nuclear regions masked by DAPI staining and cytoplasmic regions masked by Phalloidin staining, excluding areas overlapping with DAPI.

### SRPK1/2 kinase activity assay

The SRPK1 Kinase Enzyme System (#VA7558, Promega) and SRPK2 Kinase Enzyme System (#VA7561, Promega) were utilised with the ADP-Glo™ Kinase Assay (#V6930, Promega), following the manufacturer’s protocol. Briefly, 1 µl of inhibitor, diluted in 1X kinase reaction buffer, was added to the wells of a 384-well low-volume plate and combined with 8 ng of SRPK1 enzyme. Following this, 2 µl of a substrate/ATP mixture (100 µM ATP, 0.1 µg/µl MBP) was added. The reaction was incubated at room temperature for 1 hour. After incubation, 5 µl of ADP-Glo™ Reagent was added to each well, and the plate was incubated at room temperature for 40 minutes. Subsequently, 10 µl of Kinase Detection Reagent was added, and the plate was incubated for an additional 30 minutes. Luminescence was recorded.

### Proximity Ligation Assay (PLA)

The Duolink® Proximity Ligation Assay (PLA) was used to detect protein-protein interactions in KELLY cells. Samples were fixed with 4% paraformaldehyde, permeabilised with 0.1% Triton X-100, and blocked with the Duolink® blocking solution. Primary antibodies against RBM39 (1:500, Proteintech Cat# 67420-1-Ig, RRID:AB_2882660) and SRPK1 (1:250, Abcam Cat# ab90527, RRID:AB_2050339) were applied overnight at 4°C. Species-specific PLA probes (Sigma-Aldrich) were added, followed by ligation and rolling-circle amplification according to the manufacturer’s instructions.

### DNA fibre analysis

Cells were sequentially pulse-labeled with 10 μM BrdU and 200 mM IdU for 30 minutes each. KELLY cells were spotted on SuperFrost slides, air-dried for 5 minutes, and lysed with a buffer (200 mM Tris-HCl, pH 7.5, 50 mM EDTA, 0.5% SDS) to release DNA and fixed in 75% methanol/25% acetic acid, and then treated with 2.5M HCl for 1 hour to denature the DNA. After blocking with 1% BSA in PBS-T for 1 hour. BrdU was detected by anti-BrdU antibody (1:50, Abcam Cat# ab6326, RRID:AB_305426)) followed by AF647-conjugated secondary antibody (1:100, A21247, Invitrogen). IdU was labeled with mouse anti-BrdU antibody (1:50, BD Biosciences Cat# 347580, RRID:AB_10015219) and AF488-conjugated secondary antibody (1:100, Thermo Fisher Scientific Cat# A-11001, RRID:AB_2534069). Slides were mounted and imaged using a laser-free confocal microscope, with DNA track lengths measured in Fiji V2.0 (RRID:SCR_002285).

### In vivo allograft model of neuroblastoma

Syngeneic tumours derived from TH-MYCN/ALK^F1174L^ genetically engineered mouse models (GEMMs), were implanted into immune competent 129×1/SvJ mice. Group assignment was random and treatment began when the implanted tumours reached a volume of over 200 mm³. The dosing schedule continued for 2 weeks, with daily administration for 5 days followed by 2 days off. Alectinib was dissolved in saline containing 40% PEG300, 10% DMSO, and 5% Tween, and was administered orally at a dose of 10 µL/kg. Indisulam was dissolved in saline with 3.5% DMSO and 6.5% Tween80 and was administered intravenously at a dose of 5 µL/kg. Tumour tissues were excised two hours after the final dose of each compound. Half of the tumour was fixed in 4% PFA, while the other half was snap-frozen for subsequent analysis. Investigators were blind for the analysis of the study.

### Immunoassays on tissue samples

Snap frozen tumours were lysed in 5% CHAPS buffer and western blotting was carried out for RBM39 (Proteintech Cat# 67420-1-Ig, RRID:AB_2882660), ERK1/2 (Cell Signaling Technology Cat# 4695, RRID:AB_390779), pERK1/2 (Cell Signaling Technology Cat# 9101, RRID:AB_331646), phosphoepitope SR proteins (Millipore Cat# MABE50, RRID:AB_10807429) and GAPDH (Cell Signaling Technology Cat# 2118, RRID:AB_561053). ALK Meso Scale Discovery® immunoassays were performed as previously described(50)

### Immunofluorescence on tissue sections

Tumours sections were deparaffinized and rehydrated using Xylene and a graded series of alcohol. They were then boiled in 1% citric-acid buffer for 20 minutes and allowed to cool to room temperature. The sections were blocked for 1 hour in TBS with 0.01% Triton (TBST) and 5% BSA. Afterwards, γH2AX antibody (Cell Signaling Technology Cat# 9718, RRID:AB_2118009) was applied at a 1:100 dilution overnight at 4°C. Alexa Fluor 568 goat anti-rabbit antibody (1:1000) was incubated at room temperature for 1 hour. Images were captured using a confocal microscope (LSM700, LSM T-PMT) and processed with ZEN2009 software (Black version, Zeiss).

### RNA sequencing and alternative splicing analysis

Total RNA was extracted and mRNA enriched using poly-T oligo-attached magnetic beads. Strand-specific libraries were constructed and quantified by Qubit and Bioanalyzer and sequenced on an Illumina NovaSeq 6000 platform using an S4 flow cell with 150 bp paired-end reads (Novogene, Cambridge, UK). Sequencing reads were quality-filtered and clean reads were aligned to the human reference genome (hg38) using HISAT2. Gene-level counts were generated using featureCounts, and differential expression analysis was performed using DESeq2. Resulting p-values were adjusted using the Benjamini-Hochberg method to control the false discovery rate (FDR), with significance defined as adjusted p-value ≤ 0.05 Alternative splicing analysis was conducted using rMATS (v4.3.0), which detects five splice event types: skipped exon (SE), mutually exclusive exon (MXE), alternative 5′ splice site (A5SS), alternative 3 ′ splice site (A3SS), and retained intron (RI). Differential splicing events were identified based on junction-spanning reads, with significance set at FDR < 0.05.

### Statistical Analysis

Statistical analyses were conducted using GraphPad Prism 10. For normally distributed data, a Student’s t-test was applied to two-group comparisons; one-way or two-way ANOVA for three or more groups. Specific tests are detailed in figure legends.

## Acknowledgments

We are very grateful to the technicians from the Institute of Cancer Research BSU animal center, Barbara Martins da Costa and Kevin Greenslade for supporting animal work. We thank Prof. Matthew J Fuchter from Imperial College London and Dr Lizzie Tucker from Institute of Cancer Research for providing helpful comments.

## Author contributions

Conceptualization: YM, AN, HCK Methodology: YM, EP, YX, SQ, AN

Investigation: YM, EP, BMC, YX, CJ, SQ, NZ, LC, HCK, AN

Visualization: YM, YX, CJ, SQ, AN

Funding acquisition: LC, HCK, AN

Project administration: HCK, AN

Supervision: AN, HCK, LC,

Writing – original draft: YX, LC, HCK, AN

Writing – review & editing: all authors contributed to the review and editing of the paper.

### Funding

Imperial College London and China Scholarship Council joint PhD studentship 202208310101 (YM), 201808060050 (YX). AstraZeneca/NIHR Imperial BRC Imperial College Research Fellowship (AN). Cancer Research UK Fellowship RCCCDF-Nov23/100003 (AN). Cancer Research UK Discovery Programme Grant (A28278) (EP, BMC, LC); HEFCE/RAE (LC). Ovarian Cancer Action (CW, AN). Cancer Research UK convergence Science Centre PhD studentships SEBCATP-2024/100009 (NZ), CANCTA-2022\100007 (SQ).

### Data and materials availability

The RNAseq data generated in this study have been deposited in NCBI’s Gene Expression Omnibus(51) under the accession number GSE299818.

