## Supplemental materials for "Exploiting the polypharmacology of alectinib for synergistic RNA splicing disruption with RBM39 degraders"

**Supplementary Materials**

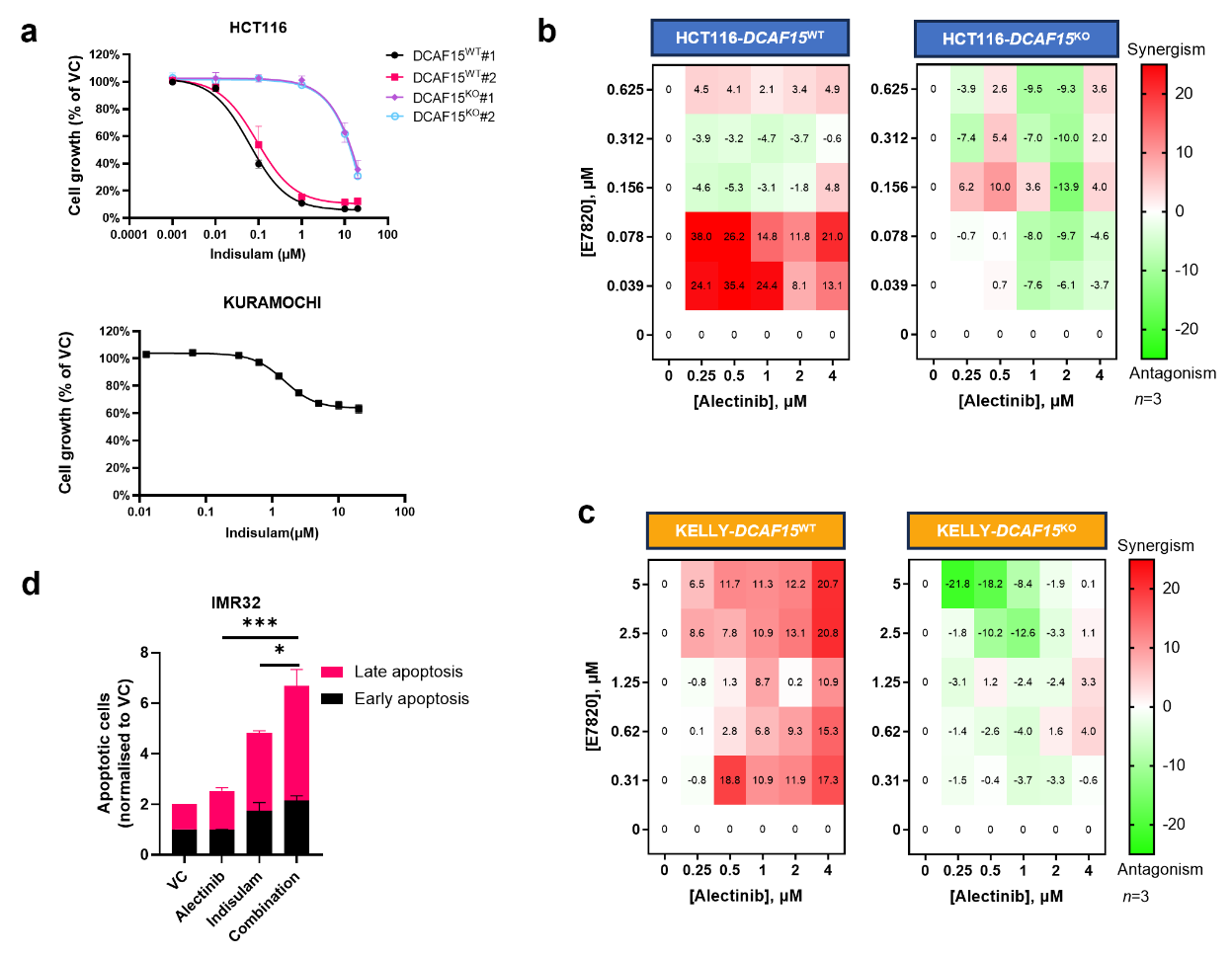

**Supplementary Fig1. Synergy of E7820 and alectinib on HCT116 and KELLY. a** Indicated cell lines were treated with indisulam for 72 hours. Cell growth was detected by SRB assay. **b-c** HCT116-*DCAF15*^WT/KO^ cell lines b and KELLY-*DCAF15*^WT/KO^ cell lines c were treated with alectinib combining E7820 for 72 hours and analysed for cell growth by SRB. Synergy scores were calculated by Synergyfinder (https://synergyfinder.fimm.fi/) using the Loewe method. **d** Apoptosis analysis via flow cytometry in IMR32 cells following exposure to 1.25µM indisulam, 4µM alectinib, or their combination for 48 hours. Cells were labelled Annexin V and PI. The significance of difference of total apoptotic cells was calculated by one-way ANOVA.

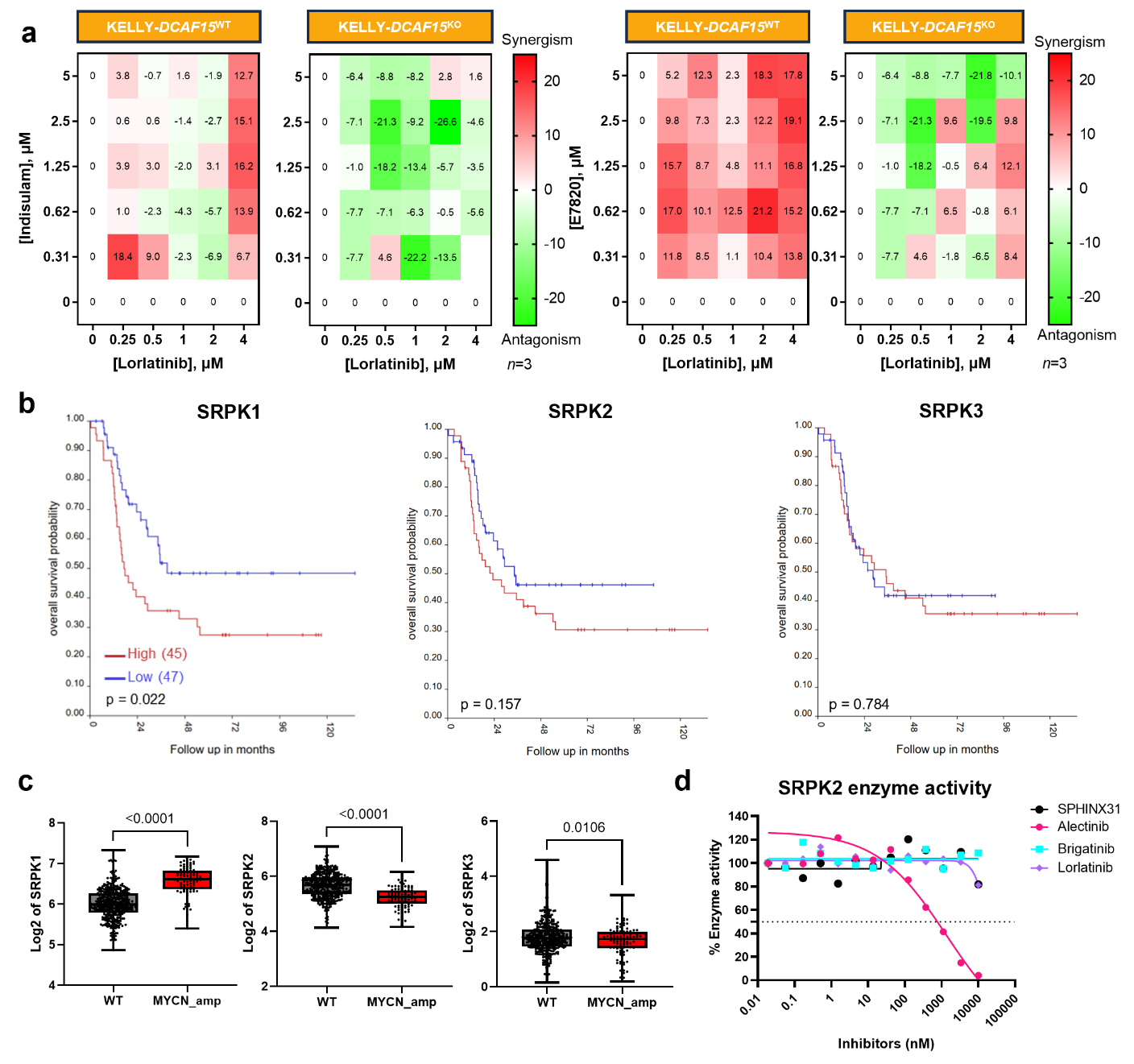
**Supplementary Fig2. Lorlatinib, SRPK2 and SRPK3 in HRNB. a** Indicated cell lines were treated with lorlatinib and indisulam concentration matrix for 72 hours. Loewe synergy scores are plotted. **b** Prognostic associations of SRPK1/2/3 expression in MYCN amplified subpopulation in the SEQC dataset, visualized using the R2 Genomics Analysis and Visualization Platform (http://r2.amc.nl), which includes gene expression data from 498 neuroblastoma patients. P values were calculated on the R2 Platform. **c** SRPK1/2/3 mRNA expression levels and MYCN status in the SEQC dataset, visualized using the R2 Genomics Analysis and Visualization Platform (http://r2.amc.nl). **d** The ADP-Glo™ Kinase Assay was performed with recombinant SRPK2 enzyme and Native Swine Myelin Basic Protein (MBP). The enzyme activity was measured by luminescent signal generated during ATP-to-ADP conversion after indicated doses of inhibitors were added to the system.

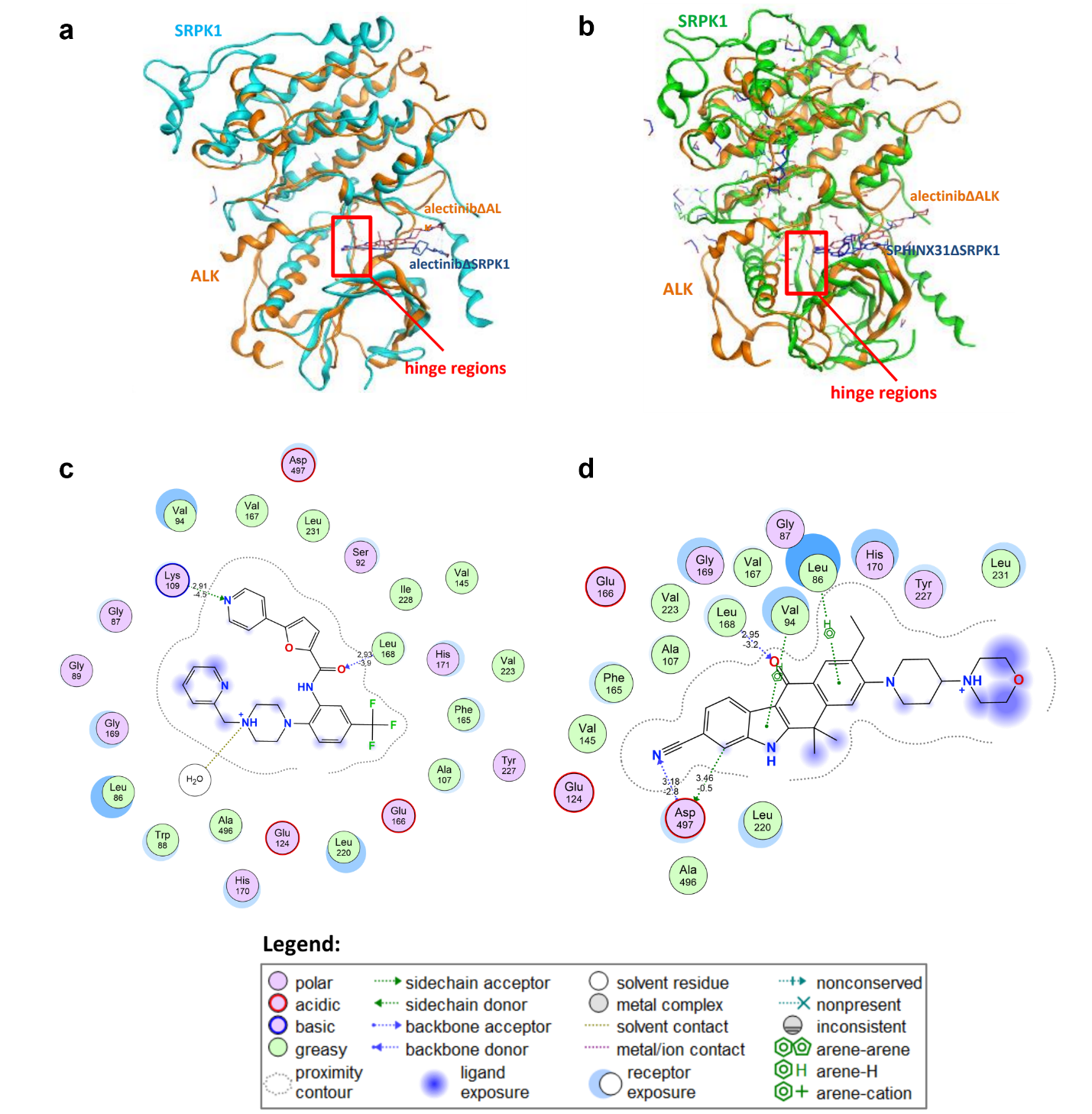

**Supplementary Fig3. Crystal Structure of the SRPK1-Alectinib Complex. a** Crystal structure of ALK complexed with alectinib (orange protein and ligand; PDB: 3AOX) superposed with crystal structure of SRPK1 complexed with alectinib (cyan protein and blue ligand; PDB: 5XV7). **b** Crystal structure of ALK complexed with alectinib (orange protein and ligand; PDB: 3AOX) superposed with crystal structure of SRPK1 complexed with SPHINX31 (green protein and blue ligand; PDB: 5XV7). **c** A map of the interactions between SPHINX31 and SRPK1 from the x-ray crystal structure (PDB: 5MY8) **d** A map of the predicted interactions between Alectinib and SRPK1 from molecular docking in 5XV7. Calculated and produced in MOE (2022.02).

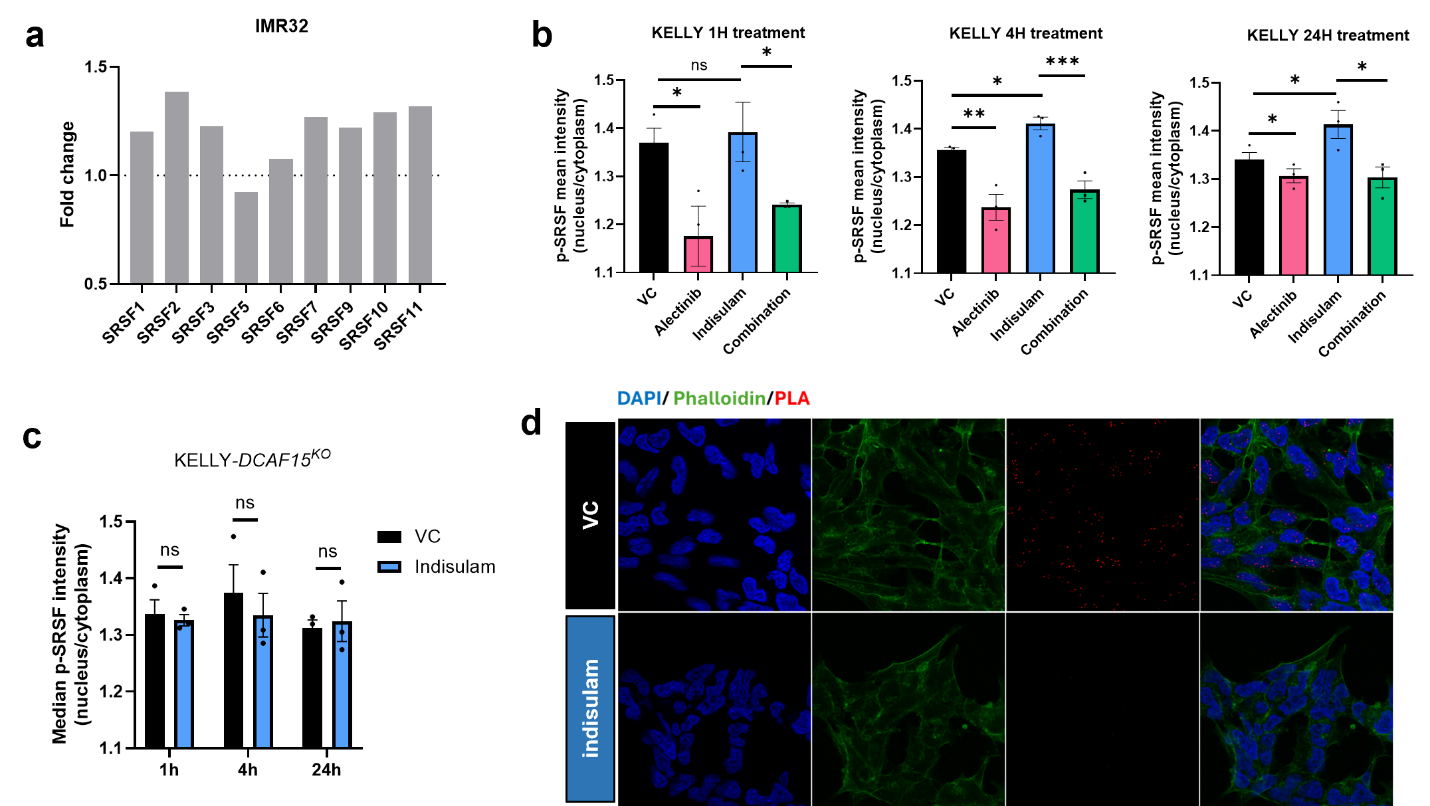

**Supplementary Fig4. SRPK1 and RBM39 interaction. a** Fold changes of SRSF family proteins after 5μM indisulam treatment in IMR32 compared to DMSO treatment by proteomics analysis. N =1. **b-c** Immunofluorescent staining of p-SRSF in KELLY-*DCAF15*^WT^ cells (**b**) or KELLY-*DCAF15*^KO^ cells (**c**), exposed to DMSO, 4μM alectinib, 1.25μM indisulam or in combination for indicated treatment time. The median values of the nuclear/cytoplasmic p-SRSF intensity from 3 independent repeats were plots. **d** Representative images of PLA assay demonstrating protein-protein interactions in KELLY cells. Nuclei are stained with DAPI (blue), actin filaments are labelled with Phalloidin (green), and PLA signals are shown in red, indicating proximity interactions between RBM39 and SRPK1. 24h treatment of 10μM indisulam was used to remove RBM39 as negative control. *p < 0.1, **p < 0.01, ***p < 0.001

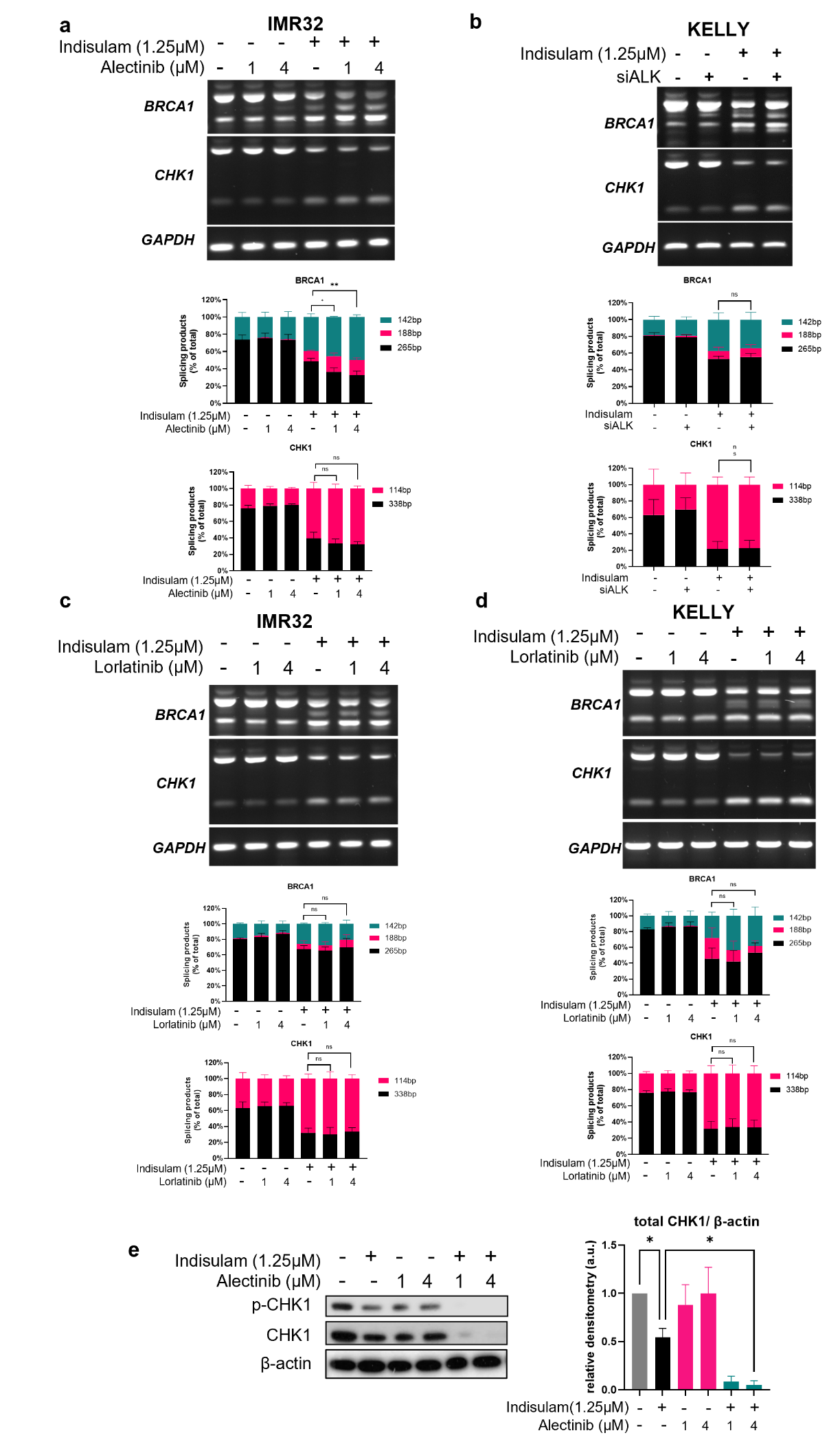

**Supplementary Fig5. ALK inhibition does not impact the splicing defect induced by indisulam. a-d** PCR validation of exon skipping events of *BRCA1* and *CHK1* in indicated cells (representative image of n=3). Cells were treated with indisulam combined with alectinib (**a**), ALK siRNA (**b**) or lorlatinib (**c-d**). *GAPDH* was included as a control. **e** Western blot of KELLY cells treated with indisulam combined alectinib at indicated doses for 48 hours (representative blot from n=3 independent experiments). n.s., no significance, * p < 0.05, **p < 0.01, ***p < 0.001.

 
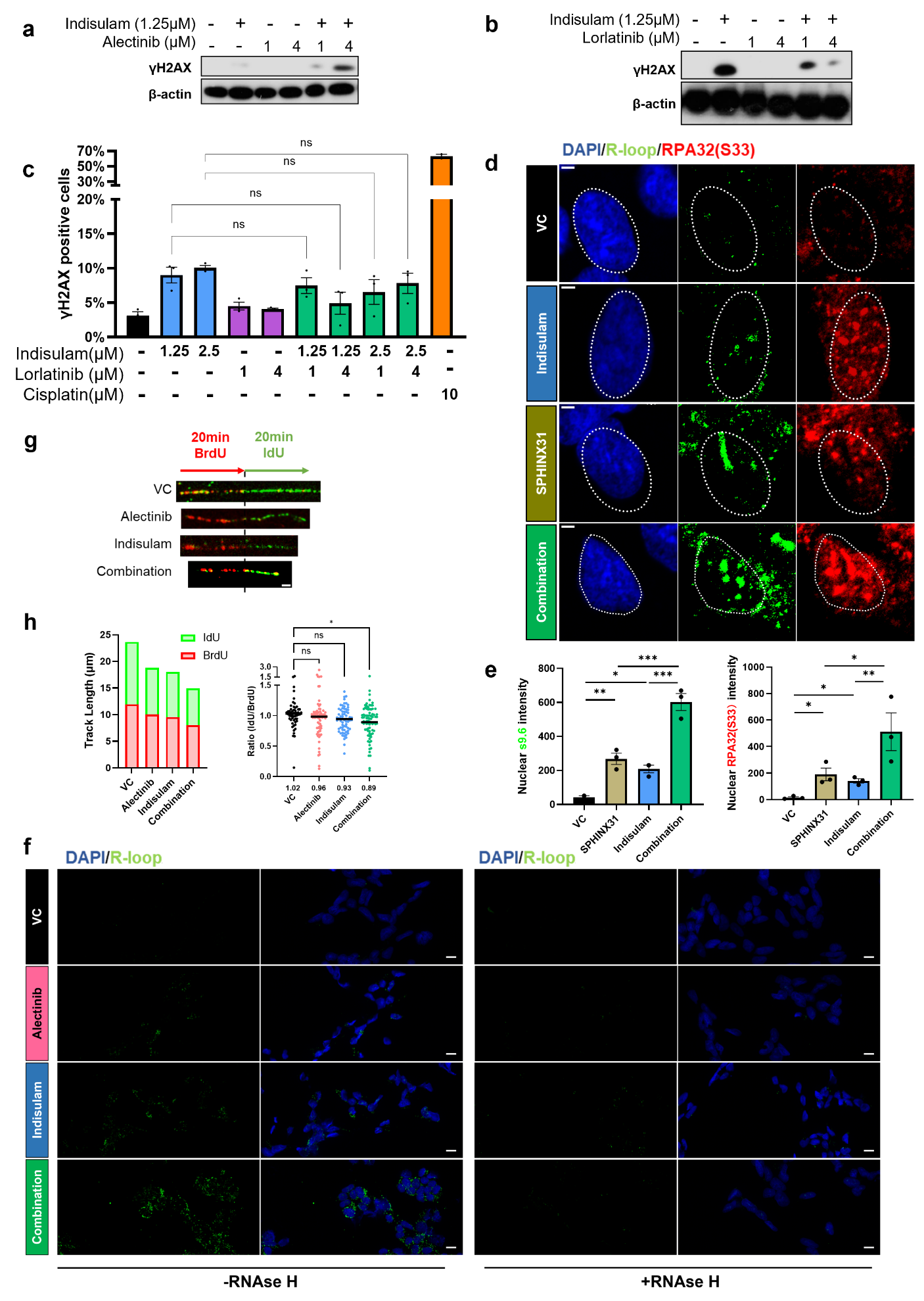

**Supplementary Fig6. ALK inhibition does not increase DNA damage induced by indisulam. a-b** Western blot and densitometry analysis of KELLY cells treated with indisulam combined alectinib a or lorlatinib b at indicated doses for 48 hours (representative blot from n=3 independent experiments). **c** Immunofluorescence staining of γH2AX in KELLY cells treated with indisulam in combination of lorlatinib. Nuclear foci >5 indicates positive cells. n=3 independent experiments. Cisplatin treatment was included as positive control. **d** Representative images for nuclear S9.6 and RPA32 (S33) foci in KELLY cells, n = 3 independent experiments, Scale bars = 2.5μm. **e** Quantification of nuclear S9.6 intensity (left) and pRPA32 intensity (right) in KELLY cells. Cells were treated with 10μM SPHINX31 monotherapy or with 1.25μM indisulam; nuclear signal intensity were measured by merging to DAPI; n>300 cells from 3 independent experiments were quantified. Significance was analysed among the median values. **f** Representative images for nuclear S9.6 staining in KELLY. Cells were treated with or without Rnase H before incubation of S9.6 antibody, Scale bars = 20μm. **g-h** Experimental scheme and representative fluorescent images for DNA fibre analysis following treatment with 4μM alectinib and 1.25μM indisulam monotherapy or in combination. Scale bar = 20 μm. At least 100 fibres were measured from each experiment. Average track length of each labelling and dot plot of BrdU/IdU ratio were shown, n = 2. n.s. = not significant, * p < 0.05, **p < 0.01, ***p < 0.001.

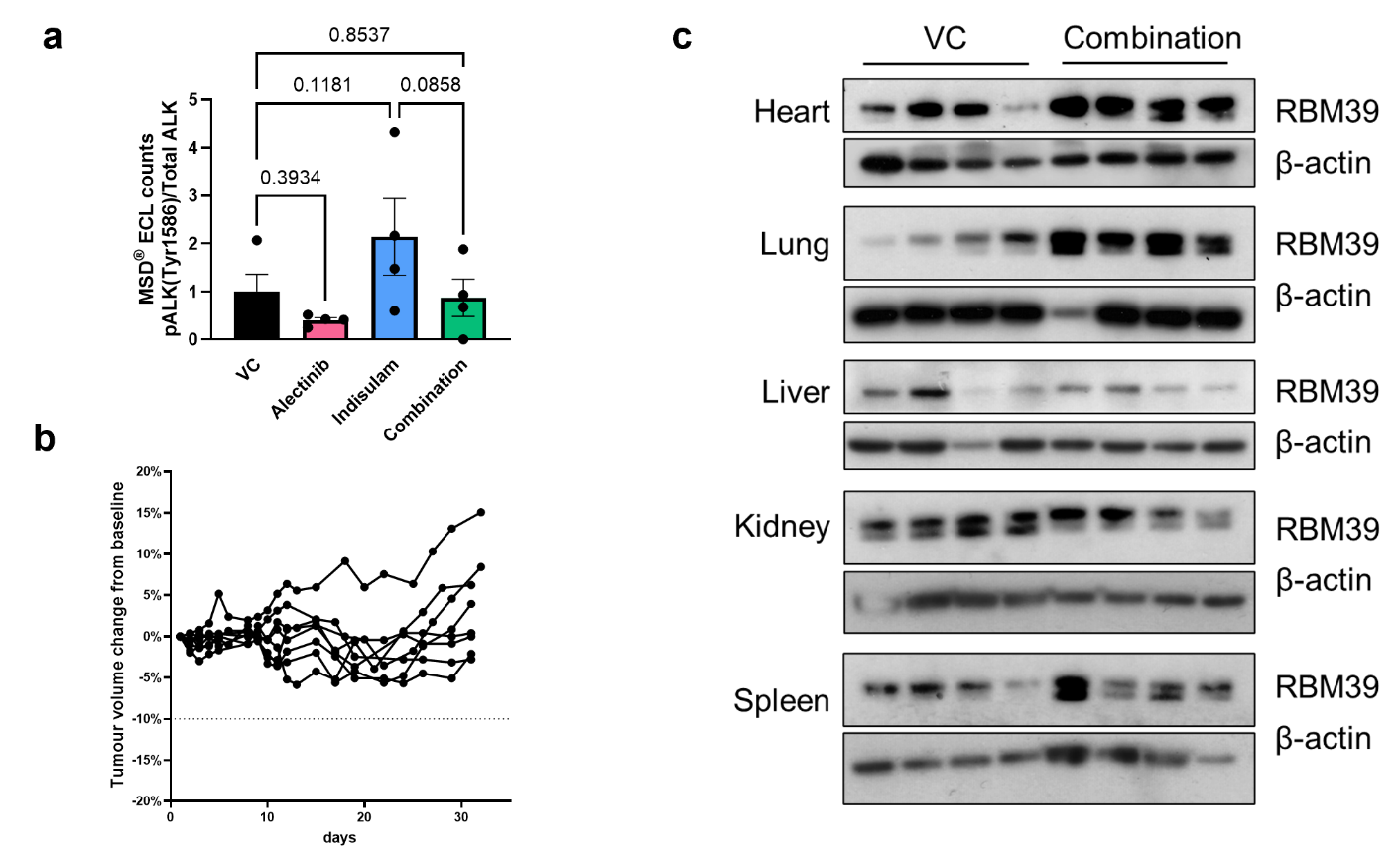

**Supplementary Fig7. Combing indisulam and alectinib shows no over toxicity. a** Protein was extracted from tumour tissues after administration of vehicle or alectinib or indisulam or combination therapy for 3 days. Phosphorylated ALK (try1586) and total ALK levels were quantified by MSD immunoassays (n = 4 per group). **b** Weight of mice receiving the combination therapy, measured daily during treatment and 2-3 times per week post-treatment. **c** Western blot assessment of RBM39 in indicated organs after treated with vehicle or combination of indisulam and alectinib for 3 days, samples were collected 2 hours post last dose (n =4).

Supplementary Table 1

| **Inhibitors** | **Targets** | **Value** | **Ref** |
| --- | --- | --- | --- |
| Alectinib | ALK* | IC50:1.9 nM | 10.1016/j.ccr.2011.04.004 |
|  | RET | IC50:4.8nM | 10.1158/1535-7163.MCT-14-0274 |
|  | SRPK1 | IC50:11 nM | 10.1016/j.chembiol.2018.01.013 |
| Lorlatinib | ALK* | Ki:<0.07 nM | 10.1021/jm500261q |
|  | ROS1* | Ki:<0.02 nM | 10.1021/jm500261q |
|  | LTK | IC50:2.7 nM | 10.1021/jm500261q |
|  | FER | IC50:3.3 nM | 10.1021/jm500261q |
| *Primary target |  |  |  |
